# Rewired NAD^+^ metabolism promotes NF-κB-mediated oxidative stress and disrupts lipid homeostasis in liver fibrosis progression

**DOI:** 10.1101/2025.08.06.668888

**Authors:** Hiu-Lok Ngan, Jacinth Wing-Sum Cheu, Kenneth Kin-Leung Kwan, Carmen Chak-Lui Wong, Hong Yan, Zongwei Cai

**Author notes:** Corresponding authors: Drs. Hong Yan and Zongwei Cai.

## Abstract

Chronic liver fibrosis significantly increases the risk of hepatocellular carcinoma (HCC), a leading cause of cancer-related deaths. However, the molecular mechanisms linking fibrosis to inflammation-associated HCC development remain unclear, complicating early diagnosis and intervention. In this study, we employ multi-omics analyses, including untargeted and targeted metabolomics, lipidomics, and transcriptomics, in a mouse model of chemically induced liver fibrosis and HCC, integrating publicly available transcriptome data from LX-2 human hepatic stellate cell (HSC) line. Our results reveal a profound rewiring of NAD^+^ metabolism as a central driver of metabolic disturbance. Analysis of bulk liver tissue shows increased activity of the kynurenine pathway of tryptophan metabolism, enhancing NAD^+^ precursor production. Hepatic nicotinamide (NAM) levels decrease due to elevated expression of NAM *N*-methyltransferase (*Nnmt*) in HSCs. Despite reduced hepatic NAM, serum NAD^+^ level rise and is compartmentalized, triggering a disruption in NAD^+^ homeostasis and activating NF-κB-mediated oxidative stress pathways. Moreover, lipid dysregulation occurs, with NF-κB dominating the regulation of SIRT1/SREBP-controlled lipogenic and cholesterogenic genes, leading to imbalances in hepatic and serum lipids. These insights elucidate connections between NAD^+^ metabolism, inflammation, and lipid dysregulation, potentially aiding in developing diagnostic biomarkers and therapeutic targets for non-viral HCC.

**Key Points:**

1. The integration of untargeted and targeted metabolomics identifies the metabolic pathways that are associated with the development of hepatocellular carcinoma (HCC) and the potential role of disturbances in NAD^+^ metabolism as a central driver of metabolic dysregulation in hepatic fibrosis, validated across transcriptomics.
2. Fibrosis-associated HCC is linked to the depletion of nicotinamide and a cascade of metabolic disturbances that promote inflammation and oxidative stress in hepatic stellate cells.
3. Targeted analysis of the metabolites in the discriminative pathways along a continuum of fibrosis-cirrhosis-HCC discovers several potential tissue and serum biomarkers for monitoring disease progression.
4. Multi-omics analysis (i.e., metabolomics, lipidomics, and transcriptomics) reveals complex interplay between NAD^+^ metabolism, NF-κB-mediated oxidative stress, and lipid homeostasis in fibrosis-associated HCC development.

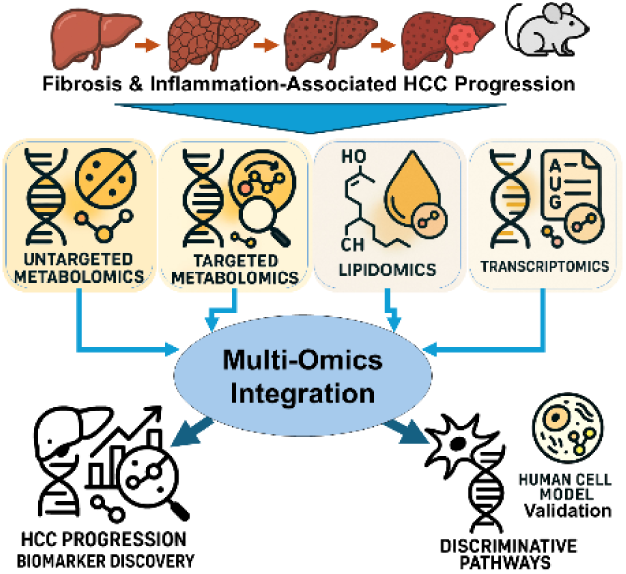

## 1. Introduction

Cancer represents a significant global health burden, according for nearly 10 million deaths worldwide in 2020 [1]. Among the various cancer types, liver cancer stands out as a particularly devastating chronic disease and is the emerging one in the United States [2]. While ranking 7^th^ in incidence, it was the 3^rd^ leading cause of cancer-related mortality [3], with a strikingly high mortality-to-incidence ratio of nearly 1:1 [3]. This underscores the lethality of liver cancer compared to other malignancies.

Early detection and intervention are critical for improving patient outcomes, yet liver cancer is often diagnosed at advanced stages, less treatable stages [4]. Hepatocellular carcinoma (HCC) is the most common subtype of liver cancer [5]. Its risk factors mostly are related to cirrhosis [6], that can be induced by hepatitis B virus (HBV) and hepatitis C virus, alcohol, and metabolism-associated factors, such as diabetes, obesity, non-alcoholic fatty liver disease (NAFLD) [7]. Chronic liver injury triggers a sustained wound-healing response characterized by the activation and differentiation of hepatic stellate cells (HSCs) into collagen-producing myofibroblasts, leading to the progressive scarring of the liver known as fibrosis [8]. Advanced liver fibrosis results in cirrhosis and involves in hepatocarcinogenesis [9]. Hence, elucidating the fibrosis-dependent mechanisms driving tumor progression will be crucial to rationalizing the development of antifibrotic therapies as an early intervention strategy.

The current body of clinical research of inflammation-associated HCC has primarily focused on analyzing serum samples from patients diagnosed with HCC [10–13], cirrhosis [13,14], fibrosis [15], or HBV infection [10,13]. Serum metabolomics is advantageous for the discovery of diagnostic biomarkers due to its relatively non-invasive sampling approach. However, in the context of early detection and disease intervention, tissue samples are necessary to examine the underlying molecular underpinnings driving the transition from chronic liver disease. Unfortunately, the availability of clinical tissue samples is often limited to the advanced stage of disease at the time of diagnosis. While there are several studies on HCC [10,13,16,17] and cirrhosis [18], tissue samples from the fibrotic stage are scarce [19]. This limitation in access to well-characterized fibrotic tissue samples represents a significant challenge that hinders our understanding of the fibrosis-dependent tumorigenic mechanisms. To fill this knowledge gap, animal models of liver fibrosis and inflammation-associated HCC are needed for a comprehensive investigation of the metabolic, signaling, and regulatory alterations in rodent liver tissue prior to clinical investigation [20,21].

In this study, we employ a diethylnitrosamine-carbon tetrachloride (DEN-CCl_4_) chemical-induced male mouse model to monitor the etiology and progression of fibrosis-associated HCC by multi-omics analyses, including inter-validation between metabolomics, lipidomics, and transcriptomics and external validation by a transcriptome set of activated human HSCs. We characterize metabolic disturbances in tryptophan metabolism, nicotinate/nicotinamide metabolism, fatty acid (FA) metabolism, alanine/aspartate/glutamate (AAG) metabolism, and citrate cycle throughout the progression stages (fibrosis, cirrhosis, and HCC) and identify several potential diagnostic serum and tissue biomarkers for disease monitoring. We confirm the dysregulated metabolic pathways discovered through our multi-omics analyses using RNA sequencing (RNA-Seq) and assess their occurrence and regulation within HSCs. Overall, our results demonstrate NAM depletion by the upregulation of NAM *N*-methyltransferase (*Nnmt*) in HSCs derives the rewiring of nicotinamide adenine dinucleotide (NAD^+^) metabolism. Disturbances in NAD^+^ homeostasis and its associated pathways trigger inflammation and lipid dysregulation, that create a pro-tumorigenic microenvironment in the fibrotic liver. These findings may ultimately contribute to improved early detection and more effective interventions to prevent the devastating progression from chronic liver disease to HCC.

## 2. Materials and Methods

### 2.1. Chemical and Reagents

Absolute Ethanol (EtOH), chloroform, formic acid (99–100%), LC/MS-grade acetonitrile (ACN), methanol (MeOH), and isopropanol (IPA) were obtained from VWR Chemicals BDH® (Poole, UK). 2-Oxo-glutaric acid, 4-aminobutanoic acid, 5-hydroxy-L-tryptophan, (S)-malic acid, chenodeoxycholic acid, cis-aconitic acid, citric acid, fumaric acid, glycocholate, indole, indoleacetic acid, isocitric acid, L-alanine, L-asparagine, L-aspartic acid, L-glutamic acid, L-glutamine, L-kynurenine, melatonin, nicotinamide (NAM), NAM adenine dinucleotide (NAD^+^), NADP^+^, NADPH, nicotinic acid, oxaloacetic acid, serine, serotonin, succinic acid, taurine, taurochenodeoxycholic acid, taurocholic acid, tryptophan, and quinolinic acid were obtained from Sigma-Aldrich (St. Louis, MO, USA). 2-Amino-4-(2-amino-3-hydroxy-phenyl)-4-oxo-butanic acid, 2-amino-4-(2-formamidophenyl)-4-oxo-butanoic acid, 5-hydroxy-indoleacetate, 5-methoxyindole-3-acetic acid, cholic acid, indole-3-acetaldehyde, maleic acid, pyridoxal, and taurochenodeoxycholic acid were purchased from Yuanye (Guangzhou, CN). 3-Hydroxyanthranilc acid, 4-pyridoxic acid, (Z)-4-amino-4-oxobut-2-enoic acid, glycochenodeoxycholic acid, indole-3-pyruvic acid, *N*-acetyl-5-hydroxytryptamine, nicotinuric acid, oxoadipic acid, and pyridoxine were collected from TargetMol (Boston, MA, USA). 5-Methoxytryptamine, 6-hydroxynicotinic acid, NADH, and *N*-methyl serotonin were obtained from Aladdin (Shanghai, CN). Nicotinic acid mononucleotide was purchased from Macklin (Shanghai, CN). 6-Hydroxy-2-oxo-1,2-dihydropyridine-3-carboxylic acid was from Bide (Hunan, CN). 2,3-Butanedione was from Alfa (Zhengzhou, CN). Total bile acid (TBA) ELISA kit was purchased from Aidisheng (Yancheng, CN). Water was purified by employing a Milli-Q® water system (Millipore, Billerica, MA, USA).

### 2.2. Animal Model for Fibrosis-Associated Hepatocellular Carcinoma

Liver cancer staging was studied by the diethylnitrosamine-carbon tetrachloride (DEN-CCl_4_) chemical-induced mouse model of fibrosis and inflammation-associated HCC since liver fibrosis was not featured in most cancer models in rodents [21]. All animal experiments and study protocols were approved by the University of Hong Kong (HKU)’s Committee on the Use of Live Animals in Teaching and Research (CULATR), in accordance with the Animals (Control of Experiments) Ordinance of Hong Kong. Mice were sacrificed if body weight loss exceeded 20%, and the maximal tumor burden allowed was not exceeded. C57BL/6J mice from the Chinese University of Hong Kong were acclimatized for 1 week at 22 ±2 °C and 60 ±5% relative humidity under a 12-hour light/dark cycle.

To develop the animal model, briefly, a single injection of a low dose of DEN (5 mg/kg) into the 2-week-old C57BL/6 male mice intraperitoneally, a well-known model of liver carcinogenesis [22]. Repeated administration of a low dose of CCl_4_ (0.5 mL/kg), a pro-fibrogenic agent, was begun at the 12^th^ week of biweekly till the 18^th^ week of age. The continuous exposure to CCl_4_ triggered the development of hepatic fibrosis and compensatory cell proliferation [23]. As the sampling plan presented in **Table A1**, liver and serum samples of 12 healthy mice, 12 fibrosis mice, 12 fibrosis age-matched control (AMC) mice, 12 cirrhosis mice, 12 cirrhosis AMC mice, 12 HCC mice, and 12 HCC AMC mice were harvested at the 18^th^ week, 20^th^ week, 20^th^ week, 28^th^ week, 28^th^ week, 47^th^ week, and 47^th^ week respectively. Liver tissue and serum samples were subsequently stored at −80 °C.

### 2.3. Preparation of Liver and Serum Extracts for Metabolomics

The whole pre-treatment was carried out at 4 °C. In brief, 25 mg of dissected mouse livers were sampled from the same region weighed and transferred into a 1.5-mL centrifuge tube. Three stainless steel beads and 1 mL of iced ice-cold 8:2 ACN/water were added to homogenize the sample by using a bullet blender homogenizer. The tissue homogenate was then centrifugated at 17,000 × g for 10 min and transferred into the new 1.5-mL centrifuge tube. Alternatively, for 100 µL of serum, 400 µL of chilled ACN was added, and the mixture was then centrifugated and transferred into the new centrifuge tube. Subsequently, the aqueous extracts were dried with an IR concentrator (NB-504CIR, N-Biotek MAX-UP, Korea). Dried samples were reconstituted by 200 µL of 1:1 MeOH/water and were mixed well thoroughly by vortex for 1 min. The pooled quality control (QC) sample was prepared by pooling 80 µL of each sample solution together and mixing. All sample solutions and QC sample were stored at –20 °C until analysis.

### 2.4. UHPLC/HRMS-Based Untargeted Metabolomics

The pre-treated tissue samples were determined by an Ultimate 3000 UHPLC system coupled with a Orbitrap Exploris 120 mass spectrometer (OE120, Thermo Fisher Scientific, San Jose, CA, USA). An ACQUITY UPLC® HSS T3 column (100 mm × 2.1 mm i.d., 1.8 µm, Waters Corporation, Manchester, UK) was used for the chromatographic separation, which was eluted at a constant flow rate of 0.300 mL/min using a gradient elution program with 0.1% of formic acid in water (A) and 0.1% of formic acid in ACN (B) as the mobile phase. The gradient elution program was as follows: 0 min, 2.0% of B; hold 1 min; 19 min, 100.0% of B; hold 2 min; back to 2% of B in 0.1 min and held for 4.9 min. The total run-time was 25 min, and the position of divert valve was set as to waste drain during the first half minute. Column temperature and autosampler temperature were set as 30 °C and 10 °C throughout the analysis, respectively. MS data was acquired by using the Xcalibur software and in positive or negative electrospray (ESI) mode separately using the optimized ionization source parameters (spray voltage: 3.0 kV for positive ESI and 2.5 kV for negative ESI; sheath gas: 50 units; auxiliary gas: 12 units; sweep gas: 0 unit; ion transfer tube temperature: 320 °C; vaporizer temperature: 350 °C). The mass spectrometer was operated in the full-scan mode from the mass-to-charge ratio (*m/z*) values of 70–1,000, and the LC/MS data was collected in profile mode at resolution of 60,000 (FWHM at *m/z* 200). Untargeted metabolomics data were pre-processed and analyzed on RStudio 2023.12.1+402 (R version 4.3.3). For level 1 identification confidence, the software *TidyMass* was employed to match the *m/z* values of molecular ion and fragments, as well as retention times (RTs) with referenced values recorded in an in-house database [24]. Level 2B annotations were achieved using the embedded external spectral library provided. The scripts can be found at https://github.com/TommyNHL/multiomics4LiverFIBHCCProj.

### 2.5. Targeted Metabolomics on the UHPLC/ESI-QqQ-MS/MS Platform

The pre-treated tissue samples were determined by an Ultimate 3000 UHPLC system coupled with a TSQ Quanitica^TM^ triple-quadrupole mass spectrometer (Thermo FisherScientific, San Jose, CA, USA). The chromatographic conditions were set as same as the conditions of untargeted metabolomics. MS data was acquired by using the Xcalibur software and in positive or negative ESI mode separately for different compound metabolites as **Table A2**. The optimized ionization source parameters include: spray voltage, 3.5 kV, static; sheath gas, 35 units; auxiliary gas, 10 units; sweep gas, 1 unit; ion transfer tube temperature, 350 °C; and vaporizer temperature, 320 °C. The mass spectrometer was run in multi-reaction monitoring (MRM) mode, and the optimal collision energy for all the selected compounds was screened from 5 to 45 V as the MRM transitions settings shown in **Table A2**.

### 2.6. LC/MS-Based Lipidomics

Similar to metabolomics, 25 mg of dissected organs were homogenized and centrifugated in 1 mL of iced ice-cold 8:2 ACN/water. Alternatively, for 100 µL of serum, 400 µL of chilled ACN was mixed and centrifugated. Lipids in the mixture were extracted by an addition of 450 µL chloroform and 150 µL water. After vortex for 1 min and centrifugation at 12,000 × g for 5 min at 6 °C, 450 µL of the upper aqueous layer was transferred into the new 1.5-mL centrifuge tube. Subsequently, the aqueous extracts were dried and reconstituted by 200 µL of 65:30:5 ACN/IPA/H_2_O, mixing well thoroughly by vortex for 1 min. The pooled QC sample was prepared by pooling 80 µL of each sample solution together and mixing. All sample solutions and QC sample were stored at –20 °C until analysis.

The pre-treated biological samples were analyzed by an Ultimate 3000 UHPLC system coupled with OE120. An ACQUITY UHPLC BEH C18 column (100 mm × 2.1 mm i.d., 1.7 µm, Waters Corporation, Manchester, UK) was used for the chromatographic separation, which was eluted at a constant flow rate of 0.260 mL/min using a gradient elution program with 0.1% of formic acid in 6:4 ACN/H_2_O with 10 mM ammonium formate (A) and 0.1% of formic acid in 9:1 IPA/ACN with 10 mM ammonium formate (B) as the mobile phase. The gradient elution program was as follows: 0 min, 30.0% of B; hold 1 min; 2 min, 45.0% of B; 7 min, 70.0% of B; hold 2 min; 17 min, 100.0% of B, hold 2 min, back to 30% of B in 1.0 min and held for 5.0 min. The total run-time was 25 min, yet the position of divert valve was set as to waste drain during the first half minute. The temperatures of column and autosampler were set as 55 °C and 18 °C throughout the analysis, respectively. MS data was acquired by using the Xcalibur software and in positive or negative ESI mode separately using the optimized ionization source parameters (spray voltage: 3.0 kV for positive ESI and 2.5 kV for negative ESI; sheath gas: 40 units; auxiliary gas: 10 units; sweep gas: 0 unit; ion transfer tube temperature: 320 °C; vaporizer temperature: 350 °C). The mass spectrometer was run in the ddMS^2^ mode with HCD collision energies of 25 and 30% from the *m/z* values of 100–1,500, and the LC/MS data was collected in profile mode at resolution of 60,000 for MS^1^ and 15,000 for MS^2^ (FWHM at *m/z* 200). Level 2B annotations of lipids were achieved using LipidSearch (Thermo Fisher Scientific, San Jose, CA, USA). In LipidSearch, a lipid defined as grade A indicates that all fatty acid (FA) chains belonging to a given lipid were completely identified, while grade C indicates the detection of lipid class-specific ions or FA-derived product ions. A python v.3.11 scripts were written for alignment and pooling lipids by class on Jupyter Notebook (Anaconda3). The scripts can be found at https://github.com/TommyNHL/multiomics4LiverFIBHCCProj.

### 2.7. Total Bile Acid Content

Cold-chilled absolute EtOH was added to 60 mg of mouse liver tissue in a 1.5-mL centrifuge tube. to homogenize the sample by using a bullet blender homogenizer. After vortex for 1 min and centrifugation at 12,000 × g for 10 min at 4 °C, 10 µL of the supernatant was sampled into a transparent 96-well plate. Sample in each well was mixed with the reagents provided by vendor, pre-treated and analyzed by ELISA reader following the manufacturer’s protocols provided. The concentration of TBA content is presented by nmol of TBA per gram of tissue sample (nmol/g).

### 2.8. Global Transcriptomics by RNA Sequencing

RNA sequencing (RNA-Seq) was performed for 6 animal subjects in each fibrosis, cirrhosis, HCC, and their AMCs groups. The 20 mg of non-tumor regions of mouse liver were subjected to total RNA extraction using the Trizol® reagent (Life Technologies, NZ). Each tissue sample was placed in 1 mL of the Trizol solution and homogenized.using a PRO200 homogenizer (Pro Scientific, CT, USA). The homogenization was carried out in two 20-second bursts at 30,000 rpm, with the sample kept on ice during the intermittent periods. Chloroform was added to the Trizol-tissue mixture, and the samples were centrifuged to separate the aqueous phase containing the extracted RNA. The RNA was then precipitated by the addition of IPA, and the precipitate was washed with 75% EtOH, dried, and resuspended in RNase/DNase-free water. Sample pretreatment processes to purify and fragment mRNA, synthesize first and second strand cDNA, ligate adapters, enrich DNA fragments, and short-read by 2 × 150 bp followed the high-throughput protocol of Illumina TruSeq^TM^ RNA Library Prep Kit. RNA-Seq data were pre-processed and analyzed on RStudio 2023.12.1+402 (R version 4.3.3). The scripts can be found at https://github.com/TommyNHL/multiomics4LiverFIBHCCProj.

### 2.9. Statistical Analysis and Data Visualization

Python v.3.11 on Jupyter Notebook (Anaconda3), R version 4.3.3 on RStudio 2023.12.1+402, and Microsoft Excel 2024 were used for statistical analyses and data visualization. Results were expressed as mean ± standard deviation (SD). Student’s *t*-test was used to assess statistically significant differences between the disease group and its AMC group, by which a value of *p* ≤0.10 was considered as statistically significant.

## 3. Results

### 3.1. Untargeted Metabolomics and Global Transcriptomics Identify Hepatic Fibrosis-Cirrhosis-HCC Progression-Related Metabolic Pathways

We initiated our study by untargeted multi-omics analyses, including reverse-phase liquid chromatography/mass spectrometry (RPLC/MS)-based metabolomics and RNA-Seq, to discover the main metabolic disturbances that occur during the progression from liver fibrosis to cirrhosis and eventual HCC using the DEN-CCl_4_ mouse model. At each disease stage, omics data of the diseased mice were evaluated against the data of age-matched controls (AMCs). For metabolite identification at this point, the confidence level followed the criteria of level 2B of the Metabolomics Standards Initiative [25], implying that the mass-to-charge ratio (*m/z*) values of the annotated metabolites’ molecular ions were matched with the chemical standards’ values registered in mass spectral libraries. A collection of annotated metabolites exhibited disease stage-specific significant differences and fold changes compared to their corresponding AMCs (**Figure 1A,B**), including lipids. Notably, their altered levels were not due to any systematic error during LC/MS assay, as all injections for quality control (QC) in both positive (**Figure 1A**) and negative (**Figure 1B**) electrospray ionization (ESI) mode were assessed to have tolerable relative standard derivations (RSDs) of <30% [26].

**Figure 1.**
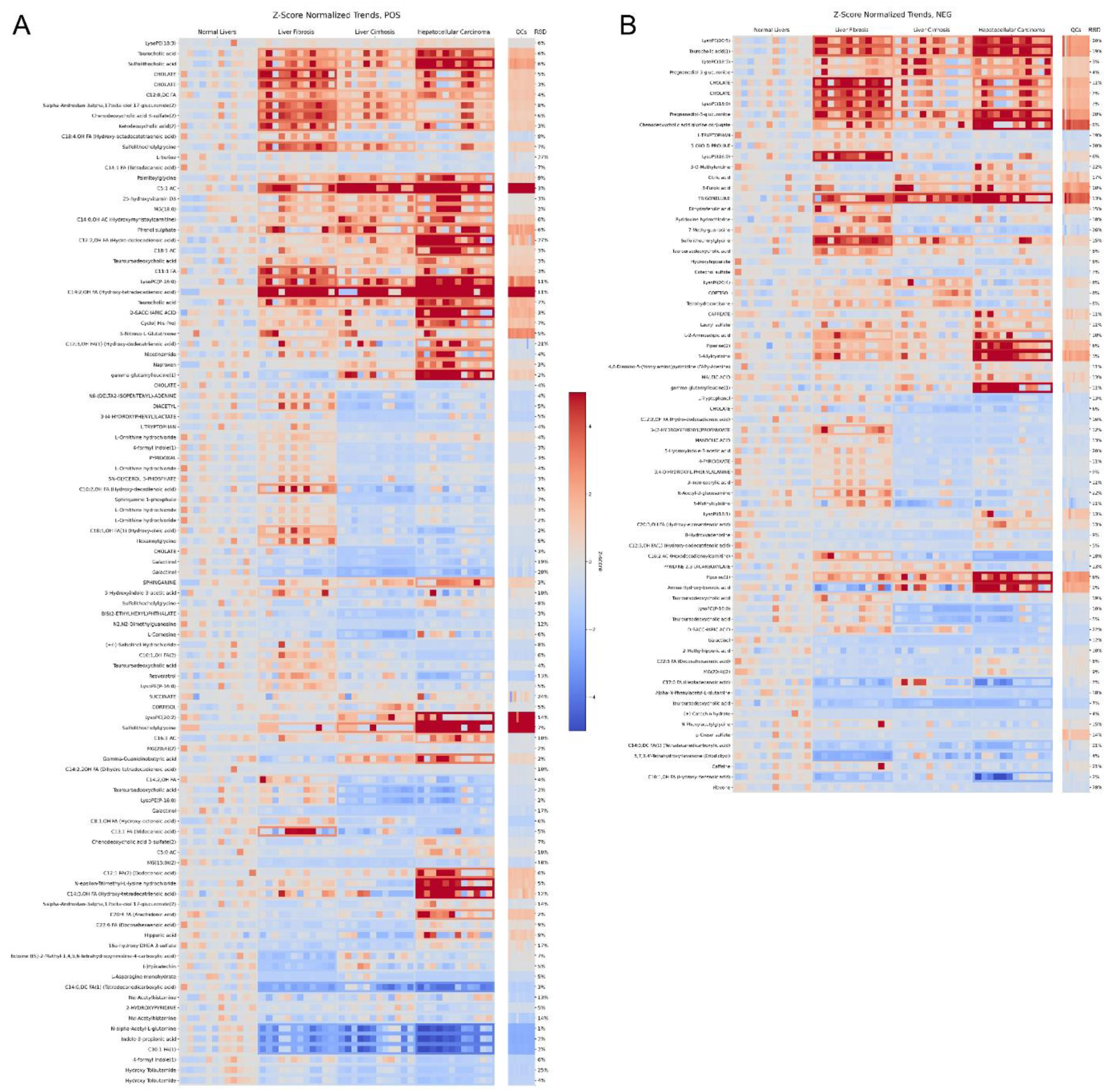
Untargeted metabolomics of the DEN-CCl_4_-treated mice liver with liver fibrosis, cirrhosis, and hepatocellular carcinoma. (A) Mass spectrometry data acquired in positive and (B) negative electrospray ionization mode. Significant metabolites from each disease stage were annotated in level 3 chemical identification confidence with a relative standard deviation (RSD) of <30% among quality control samples (QCs). Data were represented by the ratio of signal intensity of disease group to of control group, z-score normalized to normal livers.

Metabolite set enrichment analyses on the untargeted metabolomics data showed a list of pathways were significant and agreed with the pathway enrichment results of RNA-Seq (**Figure 2A,B**). Several significant pathways (i.e., AAG metabolism, tryptophan metabolism, and citrate cycle) in fibrosis (**Figure 2A,B**) were also discriminative in cirrhosis (**Figure 2C**) and HCC (**Figure 2D**). Together with nicotinate/nicotinamide metabolism (**Figure 2C,D**), they were all NAD^+^- or energy metabolism -related pathways. Targeted metabolomics was then performed to relatively quantify these compound metabolites in bulk liver tissue samples and serum, accompanied by lipidomics since FA metabolism was also enriched in global transcriptomics and relevant to energy metabolism (**Figure 2B**).

**Figure 2.**
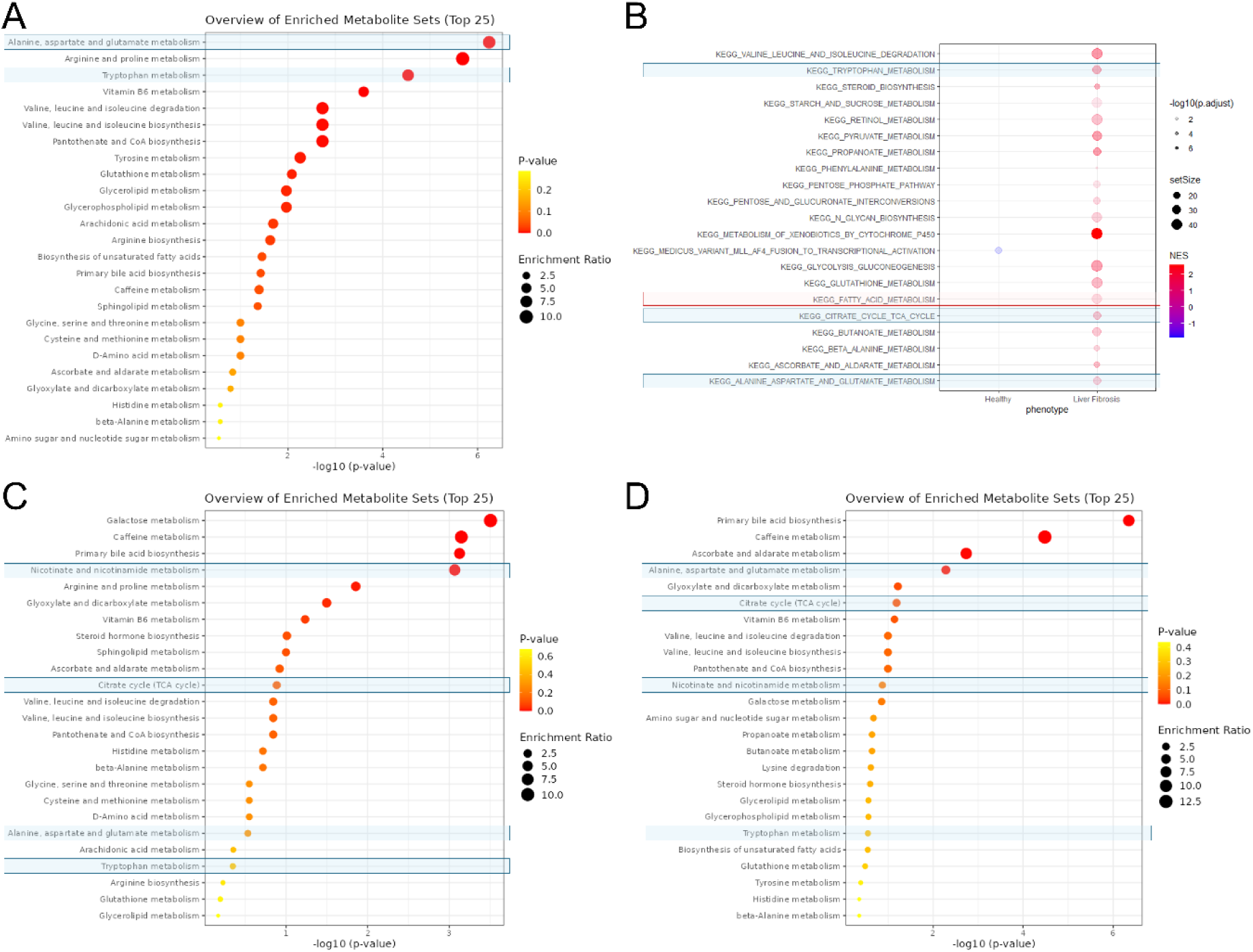
Pathway enrichment for the DEN-CCl_4_-treated mice liver with liver fibrosis, cirrhosis, and hepatocellular carcinoma. The discriminative metabolic pathways enriched from (A) metabolomics data and (B) RNA sequencing data for hepatic fibrosis, from (C) metabolomics data for cirrhotic liver, and (D) metabolomics data for carcinogenic liver. The NAD^+^-related pathways are highlighted.

### 3.2. Integration of Targeted Metabolomics and RNA Sequencing Reveals Decreased Hepatic Nicotinamide Levels Despite Upregulation of *De Novo* NAD^+^ Synthesis

To further explore the disturbances in NAD^+^ metabolism identified in the untargeted analyses, we performed targeted RPLC/tandem MS on rodent liver tissue samples. This revealed the metabolites of the kynurenine pathway of tryptophan metabolism were significantly increased in fibrotic livers (**Figure 3A,B**). Transcriptomic analysis showed that all the genes encoding regulatory enzymes were significant upregulated as differentially expressed gene (DEGs) during fibrogenesis (**Figure 3A and Table A1)**, especially the overexpression of the rate-limiting enzyme tryptophan 2,3-dioxygenase (*Tdo2*). This finding aligns with the results of tryptophan-kynurenine pathway upregulation reported by Hoshi *et al.* in their CCl_4_-induced fibrotic mouse livers [27].

**Figure 3.**
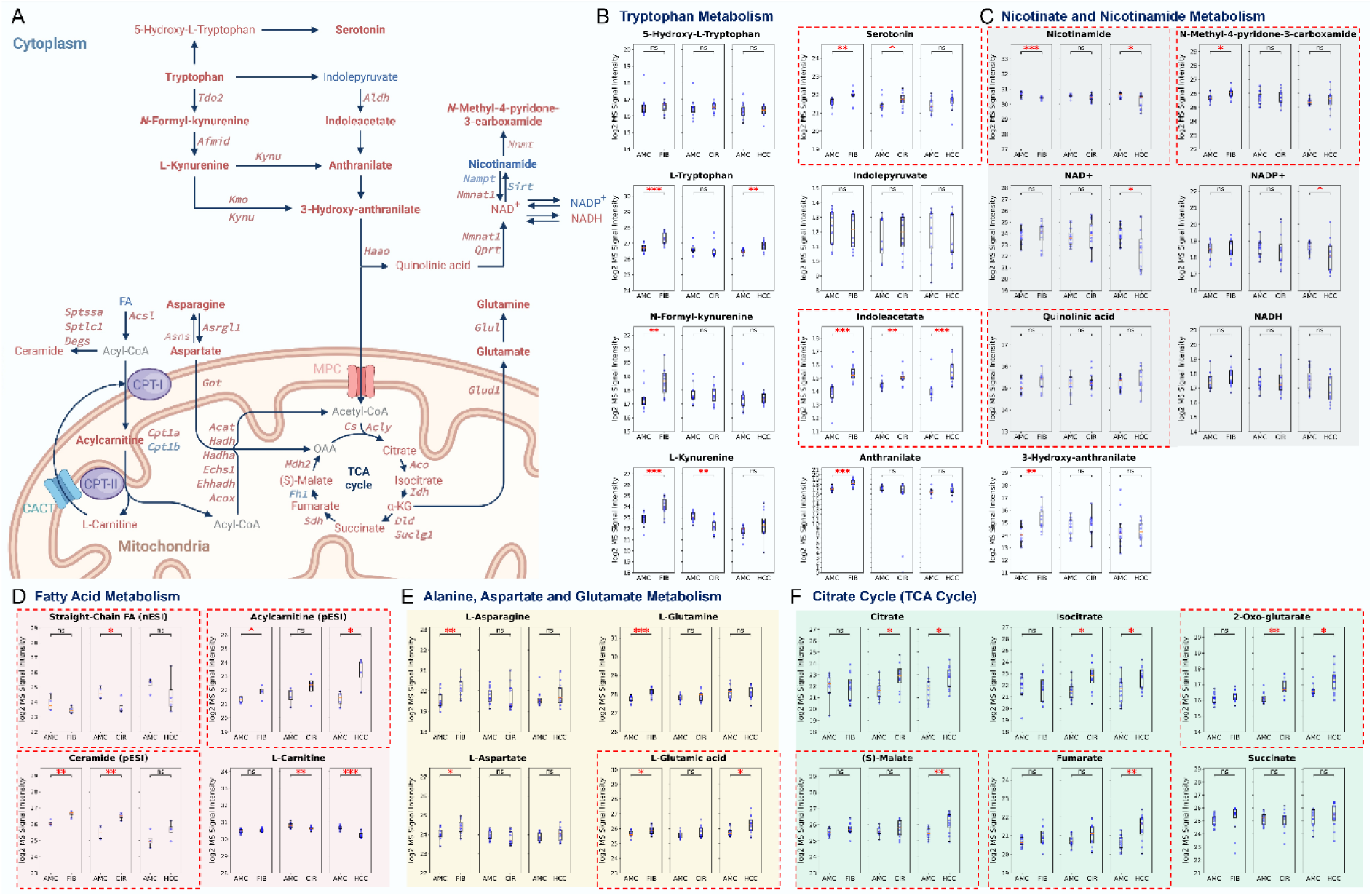
Stage-sensitive metabolomic and lipidomic changes along liver fibrosis, cirrhosis, and hepatocellular carcinoma. (A) A summary of the multi-omics data for the fibrotic mouse livers. Upregulated genes and metabolites are red-colored. Downregulated genes and metabolites are blue-colored. Non-measured metabolites are grey-colored. Bold text indicated statistical significance with p-value <0.05. Stage-sensitive alterations in metabolites and lipids in (B) tryptophan metabolism, (C) nicotinate and nicotinamide metabolism, (D) fatty acid metabolism, (E) alanine, aspartate, glutamate metabolism, and (F) citrate cycle. Potential biomarkers are dash-highlighted with red grid lines. *Abbreviations: AMC, Age-matched controls; FIB, Fibrosis; CIR, Cirrhosis; HCC, Hepatocellular carcinoma; ^, p <0.1; *, p <0.05; **, p <0.01; ***, p <0.001*.

Our targeted analysis also indicated a continuous, significant decrease in hepatic nicotinamide (NAM) levels across the fibrosis-cirrhosis-HCC disease progression (**Figure 3A,C**), despite a significant upregulation of the *de novo* NAD^+^ synthesis pathway (**Figure 3A–C)**. The RNA-Seq data for fibrosis agreed with the depletion of the available hepatic NAM pool, which was attributed to the increased expression of the NAM *N*-methyltransferase (*Nnmt*) gene **(Table A1)**, catalyzing the methylation of NAM and resulting in a significant increase of the downstream end-product *N*-methyl-4-pyridone-3-carboxamide (**Figure 3A,C**). Furthermore, we overserved downregulation of genes encoding the Sirtuin (SIRT) family of NAD^+^-dependent deacetylases, as well as the rate-limiting enzyme NAM phosphoribosyltransferase (*Nampt*) in the NAD^+^ salvage pathway. In contrast, the expression of NAM mononucleotide adenylyltransferase 1 (*Nmnat1*), which catalyzes the final step of *de novo* NAD^+^ synthesis, was upregulated. The association between the imbalance of NAM consumption/salvage and liver fibrosis via SIRT and NAD^+^ metabolism, as reported by Komatsu *et al.* [28], consolidates our findings on the nicotinate and NAM metabolism.

### 3.3. Rewiring of NAD^+^ Metabolism Dysregulates NAD^+^-Dependent Pathways

The observed disturbances in NAD^+^ homeostasis, driven by the depletion of the salvage pathway substrate NAM, prompted us to further investigate the downstream impact on NAD^+^-dependent pathways, including FA metabolism, citrate cycle, and AAG metabolism, during the fibrosis-associated HCC progression [29]. Our targeted metabolomics and lipidomics analyses revealed significant upregulation of FA beta-oxidation, as evidence by the progressive decrease of the free medium-chain saturated FA (9:0) and long-chain saturated free FAs, palmitate and stearate, and the increase in the acylcarnitine pool level (**Figure 3A,D**). The transcriptomics data indicated an increased expression of a cascade of DEGs encoding enzymes involved in FA oxidation (FAO) (**Figure 3A and Table A1)**, including the rate-limiting enzymes carnitine palmitoyltransferase 1A (*Cpt1a*), suggesting a shift towards the utilization of straight-chain free FAs as an energy source [30]. Moreover, we observed significantly increased levels of asparagine, aspartate, glutamine, and glutamate in the AAG metabolism pathway (**Figure 3A,E**) and a slightly upregulation of most citrate cycle intermediates (**Figure 3A,F**). The RNA-Seq data showed that all the related DEGs were significantly overexpressed in these metabolic pathways, corroborating the alterations observed in the targeted metabolomics data. Our findings of increased hepatic acylcarnitine and elevated TCA cycle activity were collectively aligned with the results reported by Satapati *et al.*, which demonstrated that mitochondrial citrate cycle anaplerosis was dysregulated by the upregulation of gluconeogenesis in high-fat diet male mice with obesity and hepatic insulin resistance [31].

### 3.4. Several Potential Diagnostic Tissue and Serum Metabolite Biomarkers Are Suggested for Disease Monitoring or Detection

The integration of the targeted metabolomics and transcriptomics data, along with the comparative analysis of liver tissue and serum samples, allowed us to identify several potential diagnostic biomarkers that could be feasible for monitoring the progression of liver disease. Our analyses revealed the upregulation of serotonin and indoleacetate (**Figure 3B**), the depletion of NAM, and the increase of quinolinate, *N*-methyl-4-pyridone-3-carboxamide (**Figure 3C**), the consumption of straight-chain FAs, the accumulation of acylcarnitine level, ceramide level (**Figure 3D**), glutamate (**Figure 3E**), as well as the citrate cycle intermediates like 2-oxo-glutarate, malate, and fumarate (**Figure 3F**) exhibited progressive alterations along the continuum of fibrosis-cirrhosis-HCC. These tissue biomarkers thus have the potential for translational application through fine-needle aspiration biopsy assessment.

As primary bile acid (BA) biosynthesis was also one of the significant pathways in fibrosis-associate HCC progression (**Figure 2A,C,D)**, primary BAs were analyzed in rodent samples. Taurochenodeoxycholate and taurocholate showed progressive increases at each disease stage in the DEN-CCl_4_ mice **(Figure A1A)**. Among the detectable BAs, chenodeoxycholate showed significant difference from the controls in the HCC stage in rodent samples but not progressively changed in the rodent samples **(Figure A1A,B)**. The total BA content showed no significant changes at each stage **(Figure A2)**. Serum BAs also showed no predictable pattern along the fibrosis-associated HCC progression. Chenodeoxycholate, cholate, glycochenodeoxycholate, and glycocholate only indicated stage-specific significances **(Figure A1C)**. Despite the lack of discovery of novel serum BA biomarkers, our result of glycocholate increment was in agreement with the previously reported data from another group [32].

Besides serum BAs, we identified potential serum metabolite biomarkers with progressive changes that could provide a relatively non-invasive approach for early diagnosis (**Figure 4A–C)**, such as the decreased levels of 5-hydroxytryptophan and serotonin (**Figure 4B**), and the elevated level of quinolinate (**Figure 4C**). High level of serum quinolinate has been found in liver cirrhosis patients with systematic inflammation [33]. Tryptophan, which was proposed as a potential serum biomarker by previous reports on the perturbations of tryptophan metabolism during hepatic fibrosis progression in rodent models [32], was altered with progressive increase along fibrosis-cirrhosis-HCC staging, if its relative intensity to the AMC at each disease stage is neglected.

**Figure 4.**
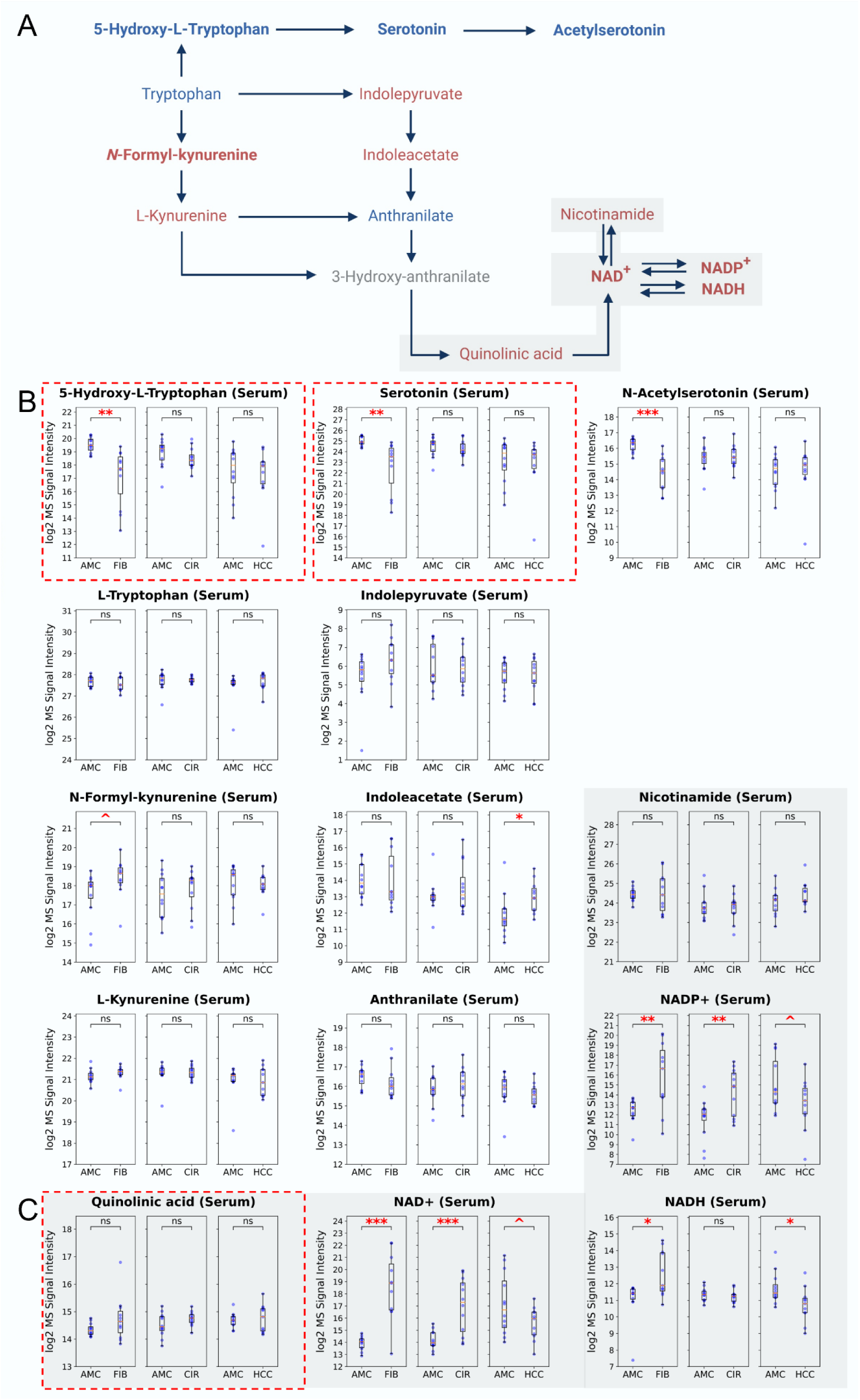
Stage-sensitive metabolomic changes in serum along liver fibrosis, cirrhosis, and hepatocellular carcinoma. (A) A summary of the metabolites in tryptophan-kynurenine pathway and NAD^+^ metabolism in the fibrotic mice serum. Upregulated metabolites are red-colored. Downregulated metabolites are blue-colored. Non-measured metabolites are grey-colored. Bold text indicated statistical significance with a p-value of <0.05. Stage-sensitive alterations in metabolites and lipids in (B) tryptophan metabolism, and (C) nicotinate and nicotinamide metabolism. Potential biomarkers are dash-highlighted with red grid lines. *Abbreviations: AMC, Age-matched controls; FIB, Fibrosis; CIR, Cirrhosis; HCC, Hepatocellular carcinoma; ^, p <0.1; *, p <0.05; **, p <0.01; ***, p <0.001*.

### 3.5. Disturbances in NAD^+^ Homeostasis and Its Associated Pathways Trigger Pro-Inflammatory Consequences

The disturbances in NAD^+^ homeostasis and the associated dysregulation of NAD^+^-dependent pathways during liver disease progression prompted us to further investigate the potential pro-inflammatory consequences using data from the transforming growth factor beta 1 (TGF-β1)-activated human HSC cell line LX-2 cell [34]. This is because the activation of HSCs, which involves the transformation of quiescent cells into proliferative, matrix-producing myofibroblasts, is recognised as a pivotal mechanism driving fibrosis in both experimental and clinical liver disease models [35]. Our transcriptomic analysis of the bulk rodent liver samples and TGF-β1-stimulated LX-2 cells revealed that the upregulation of tryptophan metabolism and serotonin synthesis may not be located within the HSCs, or the expression patterns may be temporally dissociated (**Figure 5A**). The NAD^+^ metabolism dysregulation appears to occur at an earlier stage, while the tryptophan pathway upregulation could be a later adaptive response during liver fibrogenesis. In the activated LX-2 cells, the overexpression of the DEGs, including 3-hydroxyanthranilate (3-HAA) 3,4-dioxygenase (*Haao*), quinolinate phosphoribosyl transferase (*Qprt*), *Cd38*, and nuclear factor kappa B subunit 1 (*Nfkb1*), was consistent with the RNA-Seq data from our *in vivo* studies (**Figure 5A,B,D,F and Table A1)**. Also, the downregulated genes in the *in vitro* studies, such as the AMPK regulatory subunit 5’-AMP-activated protein kinase subunit gamma-2 (*Prkag2*), *Nnmt*, *Sirt1*, and aldehyde oxidase 1 (*Aox1*), were also identified as the DEGs in the rodent fibrotic liver samples (**Figure 5A–C,E and Table A1)**.

**Figure 5.**
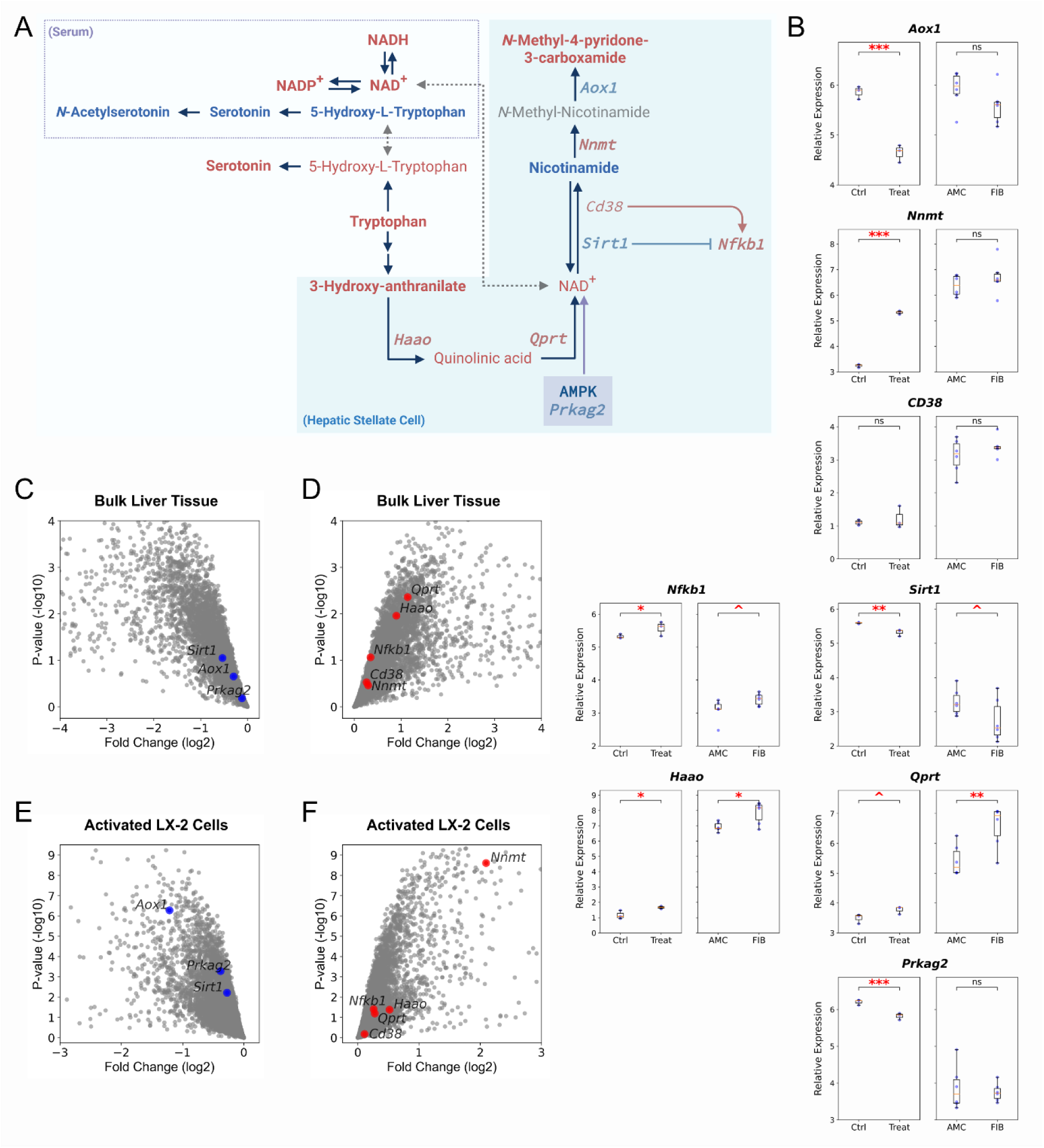
NAD^+^ metabolism and *de novo* NAD^+^ biosynthesis pathway in hepatic stellate cells. (A) A summary of the genes in *de novo* NAD^+^ biosynthesis pathway and NAD^+^ metabolism that aligned with the changes with the *in vivo* studies. Upregulated genes are red-colored. Downregulated genes are blue-colored. Bold text indicated statistical significance with a p-value of <0.05. *In vitro* and *in vivo* differentially expressed genes were represented (B) by box plots and (C–F) by volcano plots. *Abbreviations: Ctrl, Experimental control; Treat, TGF-β1-activated human HSC cell line LX-2 cells; AMC, Age-matched controls; FIB, Fibrosis; ^, p <0.1; *, p <0.05; **, p <0.01; ***, p <0.001*.

In the HSCs, the overexpression of *Nnmt*, which catalyzes the methylation of NAM, led to the accumulation of *N*1-methyl NAM. This NAM depletion was further exacerbated by the downregulation of *Aox1* (**Figure 5A–C,E and Table A1)**. The decreased NAM availability was intensified by the downregulation of *Sirt1* and AMPK/PRKAG2, leading to impaired NAM salvage. The reduced *Sirt1* activity also diminished the silencing of pro-inflammatory NF-κB signaling. Together with the upregulation of CD38, the inactivation of AMPK promoted the activation of pro-inflammatory pathways. Notably, PRKAG2 variants have been associated with liver cirrhosis [36], and 3-HAA has been reported to regulate inflammasome activation [37], further supporting our collective findings.

### 3.6. Downregulated AMPK Signaling Pathway Triggers Lipid Homeostasis Imbalance

Our metabolite set enrichment analyses revealed that the sphingolipid metabolism pathway was significantly perturbed in the early stages of fibrosis and cirrhosis (**Figure 2A,C**). The differential expression of key genes involved in the ceramide biosynthetic pathway was notably upregulated (**Figure 3A,D and Table A1)**. Meanwhile, the significant differences in acylcarnitine, a marker of FAO, relative to the levels in the AMCs was also observed (**Figure 3A,D**), suggesting a strong propensity for FA catabolism. This is supported by the evidence that the presence of *Cpt1a*, the rate-limiting enzyme in beta-oxidation, is crucial for the activation of HSCs [30]. Elevated expression of *Cpt1a* was reported in the HSCs of patients with non-alcoholic steatohepatitis (NASH), along with a positive correlation with the hepatic fibrosis score [30]. Upon activation, HSCs begin to lose their intracellular FA stores. Interestingly, the expression of *Cpt1a* was downregulated in the TGF-β1-stimulated LX-2 cells after 24 hours of incubation **(Table A1)**, contrasting with its upregulation during the earlier stages of activation (from 1 to 12 hours of incubation) reported by Fondevila *et al.* [30]. Our transcriptomic analysis of the bulk rodent liver samples and *in vitro* studies revealed that the upregulation of FAO may be temporally dissociated (**Figure 6A**), occurring at an earlier stage, while the steroid biosynthesis upregulation could be a later adaptive response during liver fibrogenesis.

**Figure 6.**
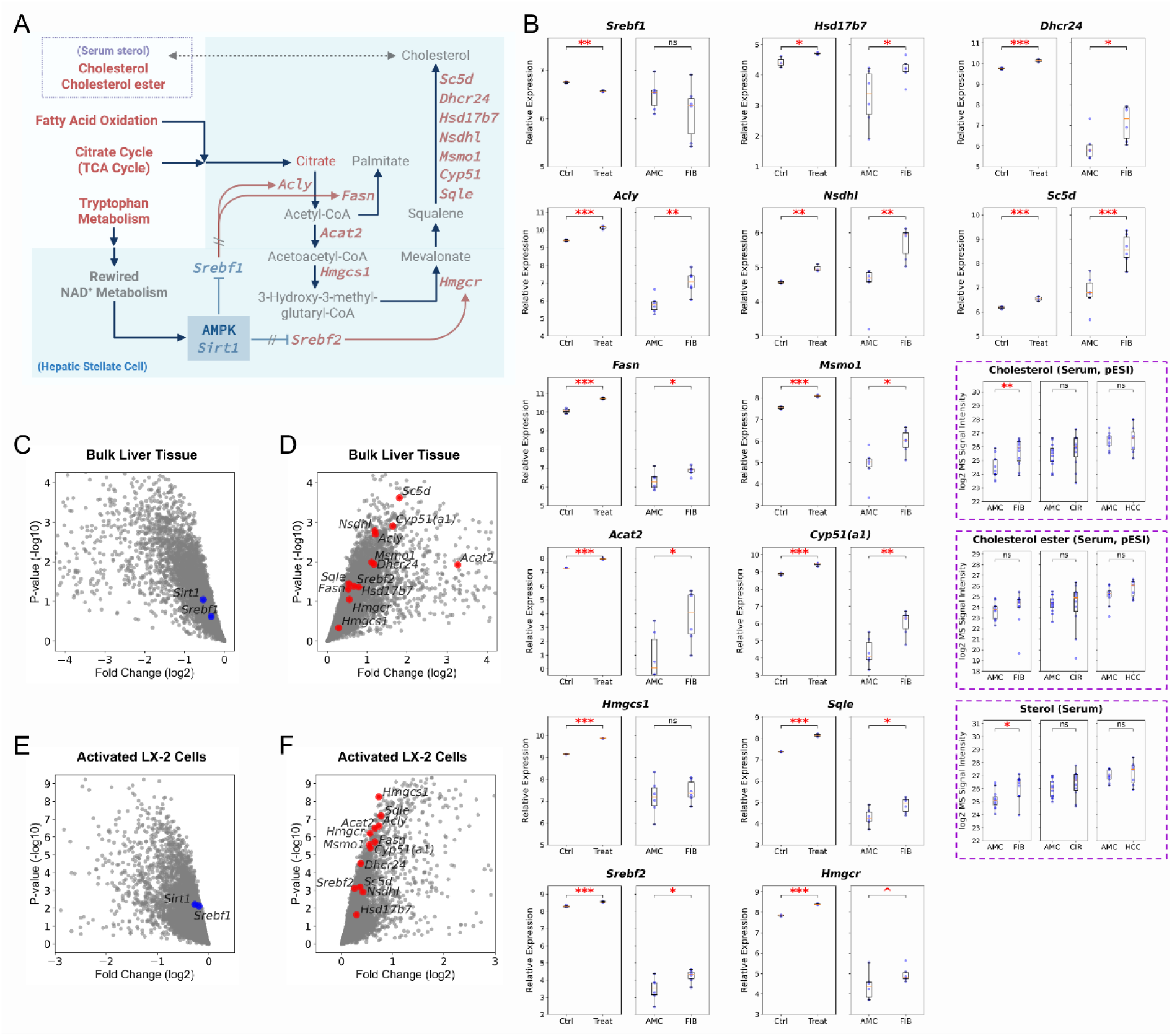
*De novo* lipogenesis and cholesterogenesis pathway in hepatic stellate cells. (A) A summary of the genes in *de novo* lipogenesis and cholesterogenesis pathway that aligned with the changes with the *in vivo* studies. Upregulated genes are red-colored. Downregulated genes are blue-colored. Bold text indicated statistical significance with a p-value of <0.05. *In vitro* and *in vivo* differentially expressed genes were represented (B) by box plots and (C–F) by volcano plots. Serum lipids are dash-highlighted with purple grid lines. *Abbreviations: Ctrl, Experimental control; Treat, TGF-β1-activated human HSC cell line LX-2 cells; AMC, Age-matched controls; FIB, Fibrosis; CIR, Cirrhosis; HCC, Hepatocellular carcinoma; ^, p <0.1; *, p <0.05; **, p <0.01; ***, p <0.001*.

In the activated HSCs, the overexpression of the DEGs, including ATP citrate lyase (*Acly*) and FA synthase (*Fasn*) in *de novo* lipogenesis (DNL), acyl-CoA:cholesterol acyltransferase 2 (*Acat2*), 3-hydroxy-3-methylglutaryl-CoA synthase 1 (*Hmgcs1*), sterol regulatory element-binding protein 2 (*Srebf2*), and a cascade of genes in the mevalonate pathway of cholesterogenesis, including 3-Hydroxy-3-methylglutaryl-CoA reductase (*Hmgcr*), was consistent with the RNA-Seq data from our *in vivo* studies (**Figure 6A,B,D,F and Table A1)**. Both *in vivo* and *in vitro* studies showed the downregulations of the AMPK-SIRT1 axis and the consequent derepression of the sterol regulatory element-binding proteins (SREBPs), which are key transcriptional regulators of DNL and cholesterogenesis in HSCs (**Figure 6A–C,E and Table A1)** [38]. This resulted in the overexpression of genes *Acly* and *Fasn*, as well as those involved in cholesterol biosynthesis, in line with the fact that lipogenesis required the overexpression of the rate-limiting enzyme *Fasn* in mouse models of obesity and NAFLD [39].

## 4. Discussion

In this comprehensive multi-omics study, we have revealed the major metabolic pathways and signaling networks that are dysregulated during liver fibrosis and inflammation-associated HCC progression. The integration of untargeted metabolomics, targeted metabolomics, lipidomics, and global transcriptomics has provided a holistic understanding of the metabolic rewiring that occurs in the early stages of liver fibrogenesis. The key metabolic pathways identified to be dysregulated include NAD^+^ metabolism, *de novo* NAD^+^ biosynthesis, sphingolipid synthesis, FAO, DNL, and cholesterogenesis. Global transcriptomics in our *in vivo* studies validates the metabolic alterations by DEGs expression. The DEGs are double confirmed by the external RNA-Seq data of TGF-β1-treated LX-2 cells [34].

The importance of NAD^+^ metabolism in regulating various cellular processes is well-established [29], and our study has revealed its significant dysregulation during liver disease progression. The state-specific changes in the levels of NAD^+^ and its precursor, NAM, suggest a complex interplay between *de novo* NAD^+^ synthesis, NAM salvage, and NAM clearance mechanisms. In the early stages of HCC progression (i.e., advanced fibrotic liver stages in fibrosis and cirrhosis), despite the upregulation of *de novo* NAD^+^ synthesis, the hepatic NAM levels were decreased (**Figure 3A,C**). This was due to the overexpression of *Nnmt* (**Figure 5A,B,D,F and Table A1)**, catalyzing the methylation of NAM. The overexpression of *Nnmt* was found during the progression of various cancers [40]. Its correlation to the cancer-associated fibroblast was reported [41], yet our *in vitro* analysis demonstrated a comparative overexpression of the DEGs encoding NAD^+^ metabolism in the activated HSCs, implying *Nnmt* has a central role of metabolic regulation of myofibroblast differentiation and cancer progression [41].

The accumulation of methylated NAM was exacerbated by the downregulation of *Aox1* (**Figure 5A–C,E and Table A1)**, an enzyme responsible for the clearance of this methylated metabolite. As a “methyl sink” in carcinogenesis, NAM is methylated and eventually excreted from the body [42]. Measurements of serum and urine methylated NAM was found to be significantly higher concentrations in patients with cirrhosis compared to healthy controls [43], suggesting that the impairment in NAM salvage mechanisms leads to the excessive clearance of this methylated metabolite from the body and decreased NAD^+^ salvage.

Attempted recovery of NAM and NAD^+^ was attained by the upregulation of the *de novo* NAD^+^ synthesis. It was evidenced by a study on the depressed expression of NAM salvage in mice [44]. Overexpression of *Haao* and *Qprt* were found in the HSCs (**Figure 5A,B,D,F and Table A1)**. This enzymatic upregulation led to the rapid production and consumption of quinolinate (**Figure 3A,C,4A)**, suggesting a metabolic adaption to maintain hepatic NAD^+^ levels. Upregulation of tryptophan-kynurenine pathway in liver was also reported in the consequence of hypoxia-causing manganese exposure [45]. Despite the increased hepatic NAD^+^ level observed (**Figure 1A,C**), the system was unable to compensate for the loss of NAM, as the NAD^+^ could not be efficiently converted back into NAM due to the downregulation of AMPK/PRKAG2 and SIRT1 (**Figure 5A–C,E and Table A1)**. As a result, the excess NAD^+^ was shunted into the systemic circulation. Increased serum NAD^+^ levels in adults with metabolic diseases, including obesity and NAFLD was reported, and there was no significant interaction between age, sex, drinking, smoking, and NAD^+^ for the metabolic diseases [46]. The high blood NAD^+^ level was not attributed from the non-hepatic NAD^+^ [47], suggesting the circulating NAD^+^ was contributed by the liver.

The decreased hepatic NAM levels, despite the upregulation of *de novo* NAD+ biosynthesis, contributed to the activation of pro-inflammatory signaling cascades in the HSCs through multiple mechanisms. Firstly, the downregulation of the AMPK-SIRT1 axis, resulting from the NAM depletion and NAD^+^ affluence, led to the derepression of NF-κB signaling (**Figure 5A,B,D,F and Table A1)**. While the decreased NAM levels and consequent impairment of this axis allowed for the unchecked activation of NF-κB in the HSCs. Additionally, the increased expression of CD38, being found on the surface of HSCs [48], further exacerbated the pro-inflammatory consequences. NAD^+^ affluence has been reported by selectively inhibition of CD38 ectoenzyme, accompanying with significant prevention from fibrosis [49], partially through the mechanisms of blunting the recruitment of immune cells and NF-κB signaling [50]. Several studies reported boosting hepatic NAD^+^ level improved liver injury and inflammation by nicotinic acid (NA) riboside supplementation [47], or NA supplementation by converting NAM into NAD^+^ [51], counteracting the activity of CD38 by that of NAMPT. However, targeting CD38 or upregulating NAMPT resulted to a similar effect as upregulation of tryptophan-kynurenine pathway, and might be not sufficient when AMPK-SIRT1 is still downregulated. Together, targeted interventions aimed at restoring NAM availability, modulating AMPK-SIRT1-NF-κB axis, and inhibiting CD38 activity may hold significant therapeutic potential for mitigating the inflammatory and fibrogenic processes.

Besides AMPK-SIRT1-NF-κB axis, our analysis revealed that rewiring of NAD^+^ metabolism also resulted to imbalance in hepatic lipid homeostasis during the progression of liver disease through AMPK-SIRT1-NF-κB-SREBP axis in HSCs [52]. The activity of SREBP2 was indicated that it can be regulated by 3-HAA metabolism [37]. The key transcription factors involved in this lipid metabolism dysregulation are the SREBPs, which are substrates of SIRT1 enzyme [38]. While the SREBP1a and SREBP1c isoforms (encoded by the *Srebf1* gene) are regulated by the AMPK-SIRT1 axis, the SREBP2 isoform (encoded by the *Srebf2* gene) is less susceptible to this repression [53]. We observed the upregulation of key genes involved in the SREBP-regulated pathways of DNL and cholesterol biosynthesis (**Figure 6A,B,D,F and Table A1)**. This suggests a shift towards increased synthesis of palmitate or other FAs, cholesterol, and sterol metabolites. Interestingly, despite this transcriptional upregulation, we detected decreased levels of palmitate and other FAs along the disease progression. This indicated that the overall tendency in FA metabolism was towards the increased acylcarnitine formation and subsequent FAO (**Figure 3A,D**). This could potentially be for the purpose of other lipid biosynthesis, such the observed upregulation of the sphingolipid signaling pathway (**Figure 3A and Table A1)** and ceramide biosynthesis in the fibrosis and cirrhosis stages.

The overexpression of genes encoding FAO, including the rate-limiting enzyme *Cpt1a* was observed *in vivo* (**Figure 3A and Table A1)**. Expression of *Cpt1a* showed to be induced by the FAO-derived reactive oxygen species [30]. In our *in vivo* studies, the FAO appears to be upregulated by the demands of saturated FAs, as reflected by the lipidomics data (**Figure 3A,D**). Although our *in vitro* data analysis showed the activated HSCs might have a delayed response in FAO, elevated expression of *Cpt1a* is a hallmark in HSC activation in the pathogenesis of hepatic fibrosis in patients with NASH, in rodents, and in TGF-β1-stimulated LX-2 cells incubated with saturated FAs-rich mediation [30]. A study also showed that killer T cells-derived interferon gamma (IFN-γ) induces activation of FAO and *Cpt1a* upregulation in cancer cells in an AMPK-dependent manner [54]. This can be used to explain why *Cpt1a* is observed *in vivo* but not in *in vitro* study, highlighting the importance of the complex liver microenvironment in shaping the metabolic adaptations.

Although increasing saturated FA (palmitate and stearate) levels by modulating NAD^+^ metabolism by NAM mononucleotide or NAM riboside supplementation have shown to ameliorate hepatic metabolic disturbances in mice with NAFLD [55], potential therapeutic strategies to retore lipid homeostasis and mitigate the fibrogenic response should targeted the AMPK-SIRT1-SREBP axis. Approaches that aim to activate AMPK have shown promise in preventing lipogenesis and NAFLD [56]. However, the role of NF-κB in the liver is complex, and its inhibition may have both beneficial and detrimental effects, warranting a cautious approach [57].

This comprehensive multi-omics study has revealed the complex dysregulation of NAD^+^ metabolism and its profound impact on the metabolic and signaling networks during the progression of liver fibrosis and inflammation-associated HCC. The depletion of NAM levels, despite the upregulation of *de novo* NAD^+^ biosynthesis, appears to be a critical driver of the pathological changes. This NAM depletion disrupts the AMPK-SIRT1 axis, leading to the aberrant activation of pro-inflammatory NF-κB signaling in the HSCs. Furthermore, the rewiring of NAD^+^ metabolism also contributes to the imbalance in hepatic lipid homeostasis through the AMPK-SIRT1-NF-κB-SREBP axis. This study provides valuable insights into the complex metabolic reprogramming that occurs during the various stages of HCC progression, offering new avenues for the development of targeted therapeutic interventions and early diagnosis.

## Abbreviations

3-HAA: 3-Hydroxyanthranilate
AAG: Alanine/aspartate/glutamate
*Acat2*: Acyl-CoA:cholesterol acyltransferase 2
*Acly*: ATP citrate lyase
AMC: Age-matched control
*Aox1*: Aldehyde oxidase 1
BA: Bile acid
CCl4: Carbon tetrachloride
*Cpt1a*: Carnitine palmitoyltransferase 1A
DEG: Differentially expressed gene
DEN: Diethylnitrosamine<colcnt=2>
DG: Diglyceride
DNL: *De novo* lipogenesis
ESI: Electrospray ionization
FA: Fatty acid
FAO: Fatty acid oxidation
*Fasn*: Fatty acid synthase
*Haao*: 3-Hydroxyanthranilate 3,4-dioxygenase
HBV: Hepatitis B virus
HCC: Hepatocellular carcinoma
*Hmgcr*: 3-Hydroxy-3-methylglutaryl-CoA reductase
*Hmgcs1*: 3-Hydroxy-3-methylglutaryl-CoA synthase 1
HSC: Hepatic stellate cell
IFN-γ: Interferon gamma
*m/z*: Mass-to-charge ratio
MS: Mass spectrometry
NA: Nicotinic acid
NAD^+^: Nicotinamide adenine dinucleotide
NAFLD: Non-alcoholic fatty liver disease
NAM: Nicotinamide
*Nampt*: Nicotinamide phosphoribosyltransferase
NASH: Non-alcoholic steatohepatitis
*Nfkb1*: Nuclear factor kappa B subunit 1
*Nmnat1*: Nicotinamide mononucleotide adenylyltransferase 1
*Nnmt*: Nicotinamide N-methyltransferase
*Prkag2*: 5’-AMP-activated protein kinase subunit gamma-2
QC: Quality control
*Qprt*: Quinolinate phosphoribosyl transferase
RNA-Seq: RNA sequencing
RPLC: Reverse-phase liquid chromatography
*Sirt*: Sirtuin
*Srebf1*: Sterol regulatory element-binding protein 1
*Srebf2*: Sterol regulatory element-binding protein 2
SREBP: Sterol regulatory element-binding protein
*Tdo2*: Tryptophan 2,3-dioxygenase
TG: Triglyceride
TGF-β1: Transforming growth factor beta 1

## Supporting Information

Supporting Information is available from the Wiley Online Library or from the author.

## Author Contributions

**Hiu-Lok Ngan:** conceptualization, resources, data curation, writing-original draft preparation, writing - review and editing, visualization. **Jacinth Wing-Sum Cheu:** writing - review and editing, visualization. **Kenneth Kin-Leung Kwan:** writing - review and editing, visualization. **Carmen Chak-Lui Wong:** conceptualization, resources, supervision. **Hong Yan:** conceptualization, resources, writing - review and editing, supervision, project administration, funding acquisition. **Zongwei Cai:** conceptualization, resources, supervision, project administration, funding acquisition. All authors have read and agreed to the published version of the manuscript.

## Acknowledgements

1. H. Y. thanks the Start-up Grant for New Academics – YAN Hong (165520). The authors from Hong Kong Baptist University appreciate the contributions made by C.C.-L. W’s team in developing and preparing the animal models.

## Ethics Statement

All animal experiments and study protocols were approved by the University of Hong Kong (HKU)’s Committee on the Use of Live Animals in Teaching and Research (CULATR), in accordance with the Animals (Control of Experiments) Ordinance of Hong Kong.

## Data Availability Statement

All study data are included in the article and/or supporting information. Processed MS-based metabolomics and lipidomics data and RNA-Seq data are available on Zenodo (see 10.5281/zenodo.17140661).

Untargeted metabolomics, targeted metabolomics, and lipidomics are utilized to analyze tissue and serum samples from a mouse model that represents a continuum of liver fibrosis, cirrhosis, and hepatocellular carcinoma (HCC). This comprehensive approach allows us to identify potential biomarkers for inflammation-associated HCC progression and further validate the relevant discriminative metabolic pathways through transcriptomics analysis of human hepatic stellate cells, which are known to be activated during hepatic fibrosis.

