## Supplementary material for "Rewired NAD^+^ metabolism promotes NF-κB-mediated oxidative stress and disrupts lipid homeostasis in liver fibrosis progression": SI Appendix

#### TABLE OF CONTENTS

**SUPPORTING FIGURES** ________________ S-2

**Figure A1. Primary bile acids' changes along liver fibrosis, cirrhosis, and HCC** ___ S-2

**Figure A2. Total bile acid changes along liver fibrosis, cirrhosis, and HCC** ________ S-3

**SUPPORTING TABLES**________________ S-4

**Table A1. Sampling plan for the DEN-$CCl_4$-induced mouse model for fibrosis and inflammation-associated hepatocellular carcinoma** ________________ S-4

**Table A2. The optimal collision energy for the selected compound metabolites** __________ S-5

**Table A3. Summary of the differentially expressed genes in DEN-$CCl_4$ mice and TGF-β1-activated human HSC cell line LX-2 cells** ________________ S-11

### SUPPORTING FIGURES

**Figure A1**

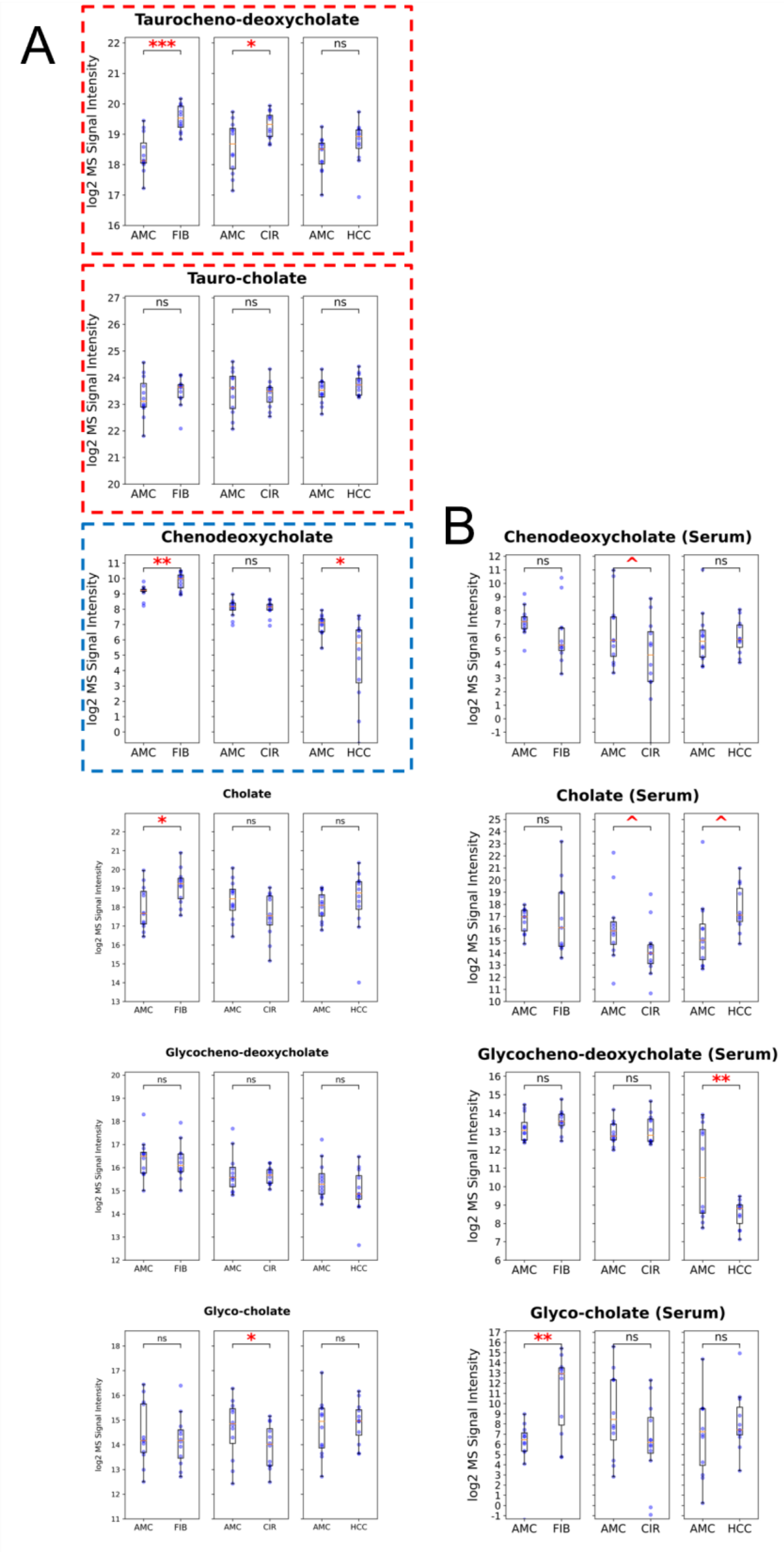


**Figure A2.Primary bile acids' changes along liver fibrosis, cirrhosis, and hepatocellular carcinoma.** (A) Bile acids in DEN-$CCl_4$ mouse livers. (B) Bile acids in DEN-$CCl_4$ mouse serum. *Abbreviations: AMC, Age-matched controls; FIB, Fibrosis; CIR, Cirrhosis; HCC, Hepatocellular carcinoma; ^, p <0.1; *, p <0.05; **, p <0.01; ***, p <0.001.*

**Figure A2**

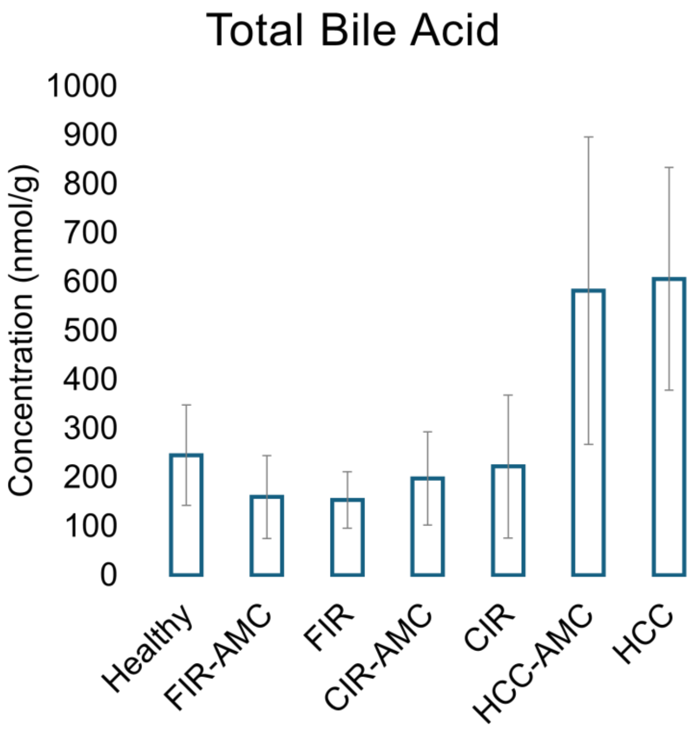


**Figure A3. Total bile acid changes along liver fibrosis, cirrhosis, and hepatocellular carcinoma.** *Abbreviations: AMC, Age-matched controls; FIB, Fibrosis; CIR, Cirrhosis; HCC, Hepatocellular carcinoma.*

### SUPPORTING TABLES

**Table A1. Sampling plan for the DEN-$CCl_4$-induced mouse model for fibrosis and inflammation-associated hepatocellular carcinoma**

| **Age** | **Status** | **Operation** |
|---|---|---|
| 02-week-old | / | DEN injection |
| 12-week-old | / | $CCl_4$ injection begun |
| 18-week-old | / | $CCl_4$ injection completed |
| 20-week-old | Fibrosis | Harvest 12 liver and serum samples |
| | Control | Harvest 12 liver and serum samples |
| 28-week-old | Cirrhosis | Harvest 12 liver and serum samples |
| | Control | Harvest 12 liver and serum samples |
| 47-week-old | HCC | Harvest 12 liver and serum samples |
| | Control | Harvest 12 liver and serum samples |

#### Table A2. The optimal collision energy for the selected compound metabolites

| Metabolites | Retention Time (min) | Polarity | Precursor Ion (*m/z*) | Product ion (*m/z*) | Collision energy (V) |
|---|---|---|---|---|---|
| Maleaic acid | 1.58 | NEG | 115.0 | 71.0 | 10 |
| | | | 115.0 | 72.0 | |
| 2-Oxo-glutaric acid | 1.45 | NEG | 145.0 | 101.0 | 10 |
| | | | 145.0 | 102.0 | |
| Quinolinic acid | 1.55 | NEG | 166.0 | 78.0 | 10 |
| | | | 166.0 | 94.0 | |
| | | | 166.0 | 122.0 | |
| 4-Pyridoxic acid | 3.46 | POS | 184.1 | 148.0 | 15 |
| | | | 184.1 | 166.1 | |
| | 3.50 | NEG | 182.0 | 108.0 | 15 |
| | | | 182.0 | 138.1 | |
| L-Tryptophan | 5.47 | POS | 205.1 | 146.1 | 10 |
| | | | 205.1 | 188.1 | |
| | 5.49 | NEG | 203.1 | 74.0 | 15 |
| | | | 203.1 | 116.0 | |
| | | | 203.1 | 142.1 | |
| | | | 203.1 | 159.1 | |
| | | | 203.1 | 166.8 | |
| Taurochenodeoxycholic acid | 10.97 | POS | 500.3 | 464.3 | 15 |
| | | | 500.3 | 482.3 | |
| | 10.98 | NEG | 498.3 | 160.8 | 25 |
| | | | 498.3 | 197.8 | |
| | | | 498.3 | 255.8 | |
| Fumaric acid | 1.58 | NEG | 115.0 | 71.0 | 10 |
| | | | 115.0 | 72.0 | |
| L-Glutamine | 0.85 | POS | 147.1 | 84.0 | 10 |
| | | | 147.1 | 130.1 | |
| | 0.85 | NEG | 145.1 | 127.1 | 10 |
| | | | 145.1 | 128.0 | |
| Pyridoxal | 0.95; 1.43 | POS | 168.1 | 94.1 | 10 |
| | | | 168.1 | 150.1 | |
| | 0.97; 1.45 | NEG | 166.1 | 108.0 | 15 |
| | | | 166.1 | 138.1 | |
| *N*-Methyl serotonin | 4.32 | POS | 191.1 | 148.1 | 15 |
| | | | 191.1 | 160.1 | |
| 5-Methoxy-indoleacetic acid | 8.64 | POS | 206.1 | 145.1 | 15 |

| | | | | | |
|---|---|---|---|---|---|
| | | | 206.1 | 160.1 | |
| Taurocholic acid | 9.92 | POS | 516.3 | 462.3 | 15 |
| | | | 516.3 | 480.3 | |
| | | | 516.3 | 498.3 | |
| L-Asparagine | 0.83 | POS | 133.1 | 74.0 | 15 |
| | 0.83 | NEG | 131.0 | 70.0 | 15 |
| | | | 131.0 | 72.0 | |
| | | | 131.0 | 95.0 | |
| | | | 131.0 | 114.0 | |
| L-Glutamic acid | 0.87 | POS | 148.1 | 84.0 | 10 |
| | | | 148.1 | 130.1 | |
| | 0.87 | NEG | 146.0 | 102.1 | 10 |
| | | | 146.0 | 128.0 | |
| Pyridoxine | 0.96; 1.59 | POS | 170.1 | 134.1 | 15 |
| | | | 170.1 | 152.1 | |
| | 0.97; 1.64 | NEG | 168.1 | 108.0 | 15 |
| | | | 168.1 | 122.1 | |
| | | | 168.1 | 138.1 | |
| | | | 168.1 | 150.1 | |
| 5-Methoxy-tryptamine | 6.19 | POS | 191.1 | 159.1 | 10 |
| | | | 191.1 | 174.1 | |
| L-Kynurenine | 4.14 | POS | 209.1 | 94.1 | 10 |
| | | | 209.1 | 136.1 | |
| | | | 209.1 | 146.1 | |
| | | | 209.1 | 192.1 | |
| | 4.17 | NEG | 207.1 | 144.0 | 10 |
| | | | 207.1 | 146.1 | |
| | | | 207.1 | 190.1 | |
| $NAD^+$ | 0.95; 1.42 | POS | 664.1 | 97.0 | 30 |
| | | | 664.1 | 136.1 | |
| | | | 664.1 | 232.1 | |
| | | | 664.1 | 428.0 | |
| | | | 664.1 | 524.1 | |
| | | | 664.1 | 542.1 | |
| | 0.95; 1.44 | NEG | 662.1 | 97.0 | 15 |
| | | | 662.1 | 134.0 | |
| | | | 662.1 | 175.0 | |
| | | | 662.1 | 346.1 | |
| | | | 662.1 | 540.1 | |
| L-Alanine | 0.85 | POS | 90.1 | 88.0 | 40 |

| | | | | | |
|---|---|---|---|---|---|
| | 0.84 | NEG | 88.0 | 71.0 | 10 |
| Succinic acid | 2.07 | NEG | 117.0 | 73.0 | 10 |
| | | | 117.0 | 99.0 | |
| L-Aspartic acid | 0.84 | POS | 134.0 | 88.0 | 10 |
| | | | 134.0 | 116.0 | |
| | 0.84 | NEG | 132.0 | 71.0 | 10 |
| | | | 132.0 | 88.0 | |
| | | | 132.0 | 115.0 | |
| 3-Hydroxy-anthranilic acid | 4.92 | POS | 154.1 | 108.0 | 10 |
| | | | 154.1 | 136.0 | |
| | 4.94 | NEG | 152.0 | 108.0 | 15 |
| | | | 152.0 | 109.0 | |
| cis-Aconitic acid | 1.98 | NEG | 173.0 | 85.0 | 10 |
| | | | 173.0 | 111.0 | |
| | | | 173.0 | 129.0 | |
| 5-Hydroxy-Indoleacetic acid | 6.23 | POS | 192.1 | 119.0 | 15 |
| | | | 192.1 | 146.1 | |
| | 6.26 | NEG | 190.1 | 144.0 | 10 |
| | | | 190.1 | 146.1 | |
| N-Acetylserotonin | 6.31 | POS | 219.1 | 160.1 | 15 |
| | | | 219.1 | 202.1 | |
| Chenodeoxycholic acid | 14.55 | NEG | 391.3 | 162.8 | 40 |
| | | | 391.3 | 197.8 | |
| NADH | 0.96; 1.66 | NEG | 664.1 | 273.0 | 30 |
| | | | 664.1 | 346.1 | |
| | | | 664.1 | 397.0 | |
| | | | 664.1 | 408.0 | |
| 4-Aminobutanoatic acid | 0.87 | POS | 104.1 | 86.1 | 10 |
| | | | 104.1 | 87.0 | |
| Nicotinamide (NAM) | 1.47 | POS | 123.1 | 80.1 | 20 |
| | | | 123.1 | 96.0 | |
| (S)-Malic acid | 1.39 | NEG | 133.0 | 71.0 | 10 |
| | | | 133.0 | 115.0 | |
| 2,6-Dihydroxy-nicotinic acid | 6.18 | POS | 174.0 | 88.0 | 15 |
| | | | 174.0 | 90.0 | |
| | | | 174.0 | 100.0 | |
| | | | 174.0 | 120.0 | |
| | | | 174.0 | 138.0 | |
| | | | 174.0 | 156.0 | |

| | | | | | |
|---|---|---|---|---|---|
| | 6.18 | NEG | 172.0 | 92.0 | 15 |
| | | | 172.0 | 93.0 | |
| Indoleacetic acid | 8.89 | POS | 176.1 | 130.1 | 15 |
| Citric acid | 1.42; 1.63 | NEG | 191.0 | 85.0 | 15 |
| | | | 191.0 | 87.0 | |
| | | | 191.0 | 111.0 | |
| 5-Hydroxy-L-Tryptophan | 4.01 | POS | 221.1 | 134.1 | 10 |
| | | | 221.1 | 162.1 | |
| | | | 221.1 | 175.1 | |
| | | | 221.1 | 204.1 | |
| | 4.08 | NEG | 219.1 | 72.0 | 15 |
| | | | 219.1 | 74.0 | |
| | | | 219.1 | 132.0 | |
| | | | 219.1 | 144.0 | |
| Cholic acid | 12.26 | NEG | 407.3 | 251.2 | 35 |
| | | | 407.3 | 289.2 | |
| | | | 407.3 | 343.3 | |
| | | | 407.3 | 363.3 | |
| $NADP^+$ | 0.97; 1.47 | POS | 744.1 | 123.1 | 40 |
| | | | 744.1 | 372.1 | |
| | | | 744.1 | 372.5 | |
| | 0.97; 1.45 | NEG | 742.1 | 79.0 | 15 |
| | | | 742.1 | 273.0 | |
| | | | 742.1 | 408.0 | |
| | | | 742.1 | 620.0 | |
| L-Serine | 0.81 | POS | 106.1 | 70.0 | 10 |
| | | | 106.1 | 88.0 | |
| | 0.81 | NEG | 104.0 | 72.0 | 10 |
| | | | 104.0 | 74.0 | |
| Nicotinic acid | 1.44 | NEG | 122.0 | 78.0 | 10 |
| | | | 122.0 | 94.0 | |
| Anthranilic acid | 7.33 | POS | 138.1 | 92.1 | 10 |
| | | | 138.1 | 120.0 | |
| | 7.34 | NEG | 136.0 | 92.0 | 15 |
| | | | 136.0 | 93.0 | |
| Serotonin | 3.96 | POS | 177.1 | 160.1 | 10 |
| DL-Isocitric acid | 1.42; 1.64 | NEG | 191.0 | 85.0 | 10 |
| | | | 191.0 | 111.0 | |
| | | | 191.0 | 173.0 | |

| | | | | | |
|---|---|---|---|---|---|
| Melatonin | 8.51 | POS | 233.1 | 174.1 | 15 |
| | | | 233.1 | 216.1 | |
| Glycochenodeoxycholic acid | 12.38 | POS | 450.3 | 76.0 | 10 |
| | | | 450.3 | 321.3 | |
| | | | 450.3 | 332.2 | |
| | | | 450.3 | 414.3 | |
| | | | 450.3 | 432.3 | |
| | | | 450.3 | 450.3 | |
| | 12.40 | NEG | 448.3 | 74.0 | 40 |
| | | | 448.3 | 384.3 | |
| | | | 448.3 | 386.3 | |
| | | | 448.3 | 404.3 | |
| Maleamic acid | 1.54 | NEG | 114.0 | 70.0 | 10 |
| | | | 114.0 | 71.0 | |
| Taurine | 0.84 | NEG | 124.0 | 80.0 | 20 |
| | | | 124.0 | 95.0 | |
| | | | 124.0 | 107.0 | |
| 6-Hydroxy-nicotinic acid | 3.20 | NEG | 138.0 | 94.0 | 15 |
| | | | 138.0 | 95.0 | |
| | | | 138.0 | 108.0 | |
| 2-Oxoadipic acid | 1.94 | NEG | 159.0 | 71.0 | 10 |
| | | | 159.0 | 115.0 | |
| Nicotinuric acid | 2.04 | POS | 181.1 | 73.1 | 15 |
| | | | 181.1 | 107.1 | |
| | | | 181.1 | 135.1 | |
| | 2.13 | NEG | 179.0 | 78.0 | 15 |
| | | | 179.0 | 108.0 | |
| | | | 179.0 | 135.1 | |
| Indolepyruvic acid | 9.74 | NEG | 202.1 | 130.1 | 10 |
| | | | 202.1 | 164.8 | |
| | | | 202.1 | 174.1 | |
| N-Formyl-kynurenine | 4.45 | POS | 237.1 | 94.1 | 15 |
| | | | 237.1 | 136.1 | |
| | | | 237.1 | 146.1 | |
| | | | 237.1 | 174.1 | |
| | | | 237.1 | 192.1 | |
| | | | 237.1 | 202.1 | |
| | | | 237.1 | 220.1 | |
| | 4.48 | NEG | 235.1 | 130.1 | 15 |
| | | | 235.1 | 146.1 | |

| | | | | | |
|---|---|---|---|---|---|
| | | | 235.1 | 174.1 | |
| | | | 235.1 | 190.1 | |
| Glycocholic acid | 10.90 | NEG | 464.3 | 160.8 | 35 |
| | | | 464.3 | 382.3 | |
| | | | 464.3 | 400.3 | |
| | | | 464.3 | 402.3 | |

**Table A3. Summary of the differentially expressed genes in DEN-$CCl_4$ mice and TGF-β1-activated human HSC cell line LX-2 cells**

| | | DEN-$CCl_4$ Mice | | | LX-2 Cells | | |
|---|---|---|---|---|---|---|---|
| Pathway | Gene | Fold Change (log2) | *P*-value | Adjusted *P* | Fold Change (log2) | *P*-value | Adjusted *P* |
| Tryptophan Metabolism | *Tph2* | / | / | / | -0.24 | 0.151 | 0.325 |
| | *Ddc* | -0.03 | 0.936 | 0.959 | / | / | / |
| | *Il4i1* | / | / | / | -0.65 | 0.070 | 0.192 |
| | *Aldh1b1* | 1.29 | 0.001 | 0.029 | 0.25 | 0.079 | 0.208 |
| | *Aldh2* | 1.72 | 0.006 | 0.053 | -0.41 | 0.000 | 0.003 |
| | *Aldh7a1* | 1.05 | 0.015 | 0.079 | -0.19 | 0.009 | 0.043 |
| | *Aldh9a1* | 0.89 | 0.022 | 0.093 | 0.37 | 0.000 | 0.002 |
| | *Aldh3a2* | 1.95 | 0.034 | 0.116 | -0.04 | 0.760 | 0.868 |
| | *Tdo2* | 1.72 | 0.000 | 0.029 | / | / | / |
| | *Ido1* | / | / | / | -0.36 | 0.368 | 0.574 |
| | *Ido2* | 0.52 | 0.140 | 0.268 | 0.10 | 0.633 | 0.787 |
| | *Afmid* | 0.73 | 0.033 | 0.115 | 0.06 | 0.546 | 0.726 |
| | *Kmo* | 1.01 | 0.011 | 0.069 | -0.01 | 0.947 | 0.973 |
| | *Kynu* | 0.67 | 0.059 | 0.158 | -0.02 | 0.845 | 0.918 |
| | *Haao* | 0.90 | 0.011 | 0.069 | 0.52 | 0.042 | 0.133 |
| Nicotinate and Nicotinamide Metabolism | *Qprt* | 1.14 | 0.004 | 0.049 | 0.28 | 0.064 | 0.179 |
| | *Nmnat1* | 0.49 | 0.063 | 0.163 | -0.07 | 0.579 | 0.749 |
| | *Nadsyn1* | -0.32 | 0.294 | 0.442 | -0.07 | 0.371 | 0.577 |
| | *Nampt* | -0.38 | 0.054 | 0.148 | 0.58 | 0.000 | 0.001 |
| | *Nmnat2* | / | / | / | -0.40 | 0.051 | 0.152 |
| | *Nmnat3* | -0.05 | 0.908 | 0.940 | / | / | / |
| | *Nadk* | -0.07 | 0.734 | 0.819 | 0.10 | 0.208 | 0.401 |
| | *Nadk2* | -0.01 | 0.981 | 0.988 | 0.19 | 0.026 | 0.095 |
| | *Nnt* | 0.75 | 0.452 | 0.596 | -0.30 | 0.004 | 0.023 |
| | *Sirt1* | -0.54 | 0.090 | 0.202 | -0.27 | 0.006 | 0.033 |
| | *Sirt2* | -0.46 | 0.023 | 0.096 | -0.02 | 0.770 | 0.874 |
| | *Sirt3* | 0.01 | 0.957 | 0.973 | -0.28 | 0.041 | 0.132 |
| | *Sirt4* | -0.69 | 0.015 | 0.080 | / | / | / |
| | *Sirt6* | -0.53 | 0.169 | 0.306 | -0.02 | 0.835 | 0.914 |
| | *Sirt7* | -0.60 | 0.010 | 0.065 | -0.10 | 0.220 | 0.416 |
| | *Sarm1* | / | / | / | 0.28 | 0.003 | 0.017 |
| | *Bst1* | / | / | / | -0.23 | 0.042 | 0.134 |

| | | | | | | | |
|---|---|---|---|---|---|---|---|
| | *Cd38* | 0.27 | 0.296 | 0.444 | 0.11 | 0.664 | 0.808 |
| | *Nnmt* | 0.30 | 0.345 | 0.494 | 2.10 | 0.000 | 0.000 |
| | *Aox1* | -0.30 | 0.223 | 0.366 | -1.21 | 0.000 | 0.000 |
| | *Aox3* | 0.29 | 0.390 | 0.537 | / | / | / |
| Citrate Cycle (TCA Cycle) | *Cs* | 0.92 | 0.012 | 0.071 | 0.14 | 0.016 | 0.067 |
| | *Aco1* | 0.60 | 0.037 | 0.121 | -0.20 | 0.017 | 0.071 |
| | *Aco2* | 0.89 | 0.037 | 0.120 | 0.04 | 0.859 | 0.927 |
| | *Idh1* | 1.24 | 0.006 | 0.053 | -0.07 | 0.344 | 0.552 |
| | *Idh2* | 1.12 | 0.008 | 0.061 | 0.01 | 0.953 | 0.976 |
| | *Idh3g* | 0.56 | 0.039 | 0.124 | 0.03 | 0.700 | 0.831 |
| | *Ogdh* | 0.29 | 0.209 | 0.352 | 0.03 | 0.681 | 0.820 |
| | *Dlst* | 2.95 | 0.117 | 0.240 | 0.01 | 0.842 | 0.917 |
| | *Dld* | 0.96 | 0.001 | 0.033 | 0.08 | 0.205 | 0.397 |
| | *Suclg2* | 2.27 | 0.128 | 0.254 | -0.56 | 0.000 | 0.001 |
| | *Suclg1* | 1.59 | 0.005 | 0.051 | 0.10 | 0.266 | 0.469 |
| | *Sucla2* | 0.59 | 0.184 | 0.322 | 0.23 | 0.458 | 0.654 |
| | *Sdhc* | 0.74 | 0.037 | 0.121 | 0.01 | 0.899 | 0.948 |
| | *Sdhd* | 1.11 | 0.003 | 0.044 | 0.02 | 0.820 | 0.905 |
| | *Sdha* | 1.28 | 0.002 | 0.038 | -0.08 | 0.120 | 0.278 |
| | *Sdhb* | 0.73 | 0.040 | 0.126 | 0.00 | 0.993 | 0.998 |
| | *Fh1* | -0.51 | 0.071 | 0.175 | 0.05 | 0.640 | 0.792 |
| | *Mdh2* | 1.29 | 0.004 | 0.045 | 0.04 | 0.520 | 0.704 |
| Fatty Acid Metabolism | *Pnpla2* | 0.56 | 0.026 | 0.102 | -0.36 | 0.002 | 0.013 |
| | *Lipe* | -1.27 | 0.008 | 0.062 | -0.44 | 0.011 | 0.052 |
| | *Mgll* | -0.73 | 0.029 | 0.107 | -0.06 | 0.608 | 0.770 |
| Fatty Acid Metabolism / Fatty Acid Degradation | *Acsl5* | 2.30 | 0.000 | 0.025 | -0.34 | 0.313 | 0.519 |
| | *Acsl3* | 0.75 | 0.002 | 0.040 | 0.77 | 0.000 | 0.000 |
| | *Acsl1* | -0.30 | 0.314 | 0.462 | -0.04 | 0.744 | 0.857 |
| | *Acsl4* | 0.13 | 0.548 | 0.673 | 0.43 | 0.001 | 0.008 |
| | *Sptssa* | 0.70 | 0.031 | 0.111 | 0.04 | 0.633 | 0.787 |
| | *Sptlc1* | 0.98 | 0.002 | 0.040 | 0.13 | 0.210 | 0.403 |
| | *Cers6* | -1.69 | 0.001 | 0.029 | -0.05 | 0.486 | 0.676 |
| | *Cers4* | -1.12 | 0.025 | 0.101 | -0.03 | 0.877 | 0.937 |
| | *Degs1* | 0.84 | 0.009 | 0.064 | 0.01 | 0.849 | 0.921 |
| | *Cpt1a* | 0.89 | 0.011 | 0.068 | -0.51 | 0.001 | 0.007 |
| | *Cpt1b* | -2.12 | 0.062 | 0.161 | -0.09 | 0.432 | 0.632 |
| | *Slc25a20* | 0.04 | 0.827 | 0.886 | -0.27 | 0.126 | 0.287 |
| | *Slc25a29* | 0.23 | 0.521 | 0.651 | -0.21 | 0.051 | 0.153 |

| | | | | | | | |
|---|---|---|---|---|---|---|---|
| | *Cpt2* | 0.06 | 0.852 | 0.903 | -0.10 | 0.318 | 0.524 |
| | *Acox1* | 1.04 | 0.005 | 0.050 | -0.02 | 0.796 | 0.891 |
| | *Acox3* | 0.15 | 0.557 | 0.680 | 0.72 | 0.000 | 0.002 |
| | *Ehhadh* | 1.60 | 0.004 | 0.045 | -0.44 | 0.044 | 0.139 |
| | *Echs1* | 1.33 | 0.002 | 0.037 | 0.03 | 0.725 | 0.847 |
| | *Hadha* | 0.79 | 0.015 | 0.080 | 0.09 | 0.337 | 0.544 |
| | *Hadh* | 0.80 | 0.020 | 0.090 | -0.33 | 0.037 | 0.122 |
| | *Acaa1* | 0.18 | 0.337 | 0.485 | -0.01 | 0.896 | 0.947 |
| | *Acaa2* | 1.44 | 0.003 | 0.044 | -0.35 | 0.008 | 0.039 |
| | *Hadhb* | 1.11 | 0.005 | 0.051 | -0.05 | 0.561 | 0.737 |
| Fatty Acid Metabolism / Mevalonate Pathway | *Acat1* | 0.77 | 0.033 | 0.113 | -0.09 | 0.323 | 0.530 |
| | *Acat2* | 3.27 | 0.012 | 0.071 | 0.65 | 0.000 | 0.000 |
| | *Acat3* | 0.72 | 0.018 | 0.085 | / | / | / |
| | *Srebf1* | -0.33 | 0.241 | 0.386 | -0.19 | 0.008 | 0.039 |
| | *Srebf2* | 0.66 | 0.041 | 0.128 | 0.26 | 0.001 | 0.007 |
| | *Acly* | 1.21 | 0.002 | 0.037 | 0.73 | 0.000 | 0.000 |
| | *Acss2* | 1.69 | 0.001 | 0.029 | 0.18 | 0.021 | 0.082 |
| | *Acss3* | -0.13 | 0.557 | 0.681 | -0.86 | 0.001 | 0.007 |
| | *Fasn* | 0.52 | 0.049 | 0.142 | 0.65 | 0.000 | 0.000 |
| | *Hmgcs1* | 0.29 | 0.462 | 0.604 | 0.73 | 0.000 | 0.000 |
| | *Hmgcr* | 0.56 | 0.089 | 0.201 | 0.55 | 0.000 | 0.000 |
| Fatty Acid Metabolism / Steroid Biosynthesis | *Mvk* | 0.00 | 0.999 | 0.999 | 0.21 | 0.063 | 0.176 |
| | *Pmvk* | 1.40 | 0.062 | 0.162 | -0.26 | 0.078 | 0.205 |
| | *Mvd* | 0.04 | 0.924 | 0.950 | 0.87 | 0.000 | 0.000 |
| | *Sqle* | 0.54 | 0.035 | 0.118 | 0.77 | 0.000 | 0.000 |
| | *Fdft1* | 0.87 | 0.015 | 0.079 | 0.21 | 0.299 | 0.504 |
| | *Lss* | 0.78 | 0.017 | 0.083 | -0.31 | 0.002 | 0.014 |
| | *Cyp51(a1)* | 1.64 | 0.001 | 0.032 | 0.56 | 0.000 | 0.000 |
| | *Tm7sf2* | 1.74 | 0.003 | 0.040 | 0.00 | 0.994 | 0.998 |
| | *Lbr* | 0.56 | 0.023 | 0.095 | -0.35 | 0.001 | 0.006 |
| | *Msmo1* | 1.11 | 0.010 | 0.067 | 0.54 | 0.000 | 0.000 |
| | *Nsdhl* | 1.19 | 0.002 | 0.035 | 0.42 | 0.001 | 0.010 |
| | *Hsd17b7* | 0.79 | 0.043 | 0.131 | 0.29 | 0.023 | 0.088 |
| | *Dhcr24* | 1.17 | 0.011 | 0.070 | 0.38 | 0.000 | 0.000 |
| | *Sc5d* | 1.80 | 0.000 | 0.024 | 0.36 | 0.001 | 0.006 |
| | *Dhcr7* | 1.12 | 0.003 | 0.043 | 0.67 | 0.000 | 0.000 |
| Alanine, Aspartate, | *Asns* | 0.23 | 0.674 | 0.775 | 0.48 | 0.000 | 0.002 |
| | *Asrgl1* | 0.60 | 0.037 | 0.121 | -0.22 | 0.195 | 0.385 |

| | | | | | | | |
|---|---|---|---|---|---|---|---|
| and | *Got1* | 0.97 | 0.050 | 0.143 | -0.28 | 0.012 | 0.055 |
| Glutamate | *Got2* | 1.34 | 0.006 | 0.056 | 0.06 | 0.388 | 0.593 |
| Metabolism | *Glud1* | 0.82 | 0.010 | 0.066 | -0.30 | 0.008 | 0.040 |
| | *Glul* | 0.76 | 0.060 | 0.159 | -0.27 | 0.006 | 0.033 |
| AMPK | *Stk11* | -0.65 | 0.007 | 0.058 | -0.21 | 0.139 | 0.306 |
| Signaling | *Prkaa2* | 0.31 | 0.147 | 0.278 | -0.76 | 0.001 | 0.009 |
| Pathway / | *Prkab1* | -0.07 | 0.743 | 0.825 | 0.00 | 0.998 | 0.999 |
| Longevity | *Prkab2* | -0.22 | 0.589 | 0.708 | 0.61 | 0.000 | 0.001 |
| Regulating | *Prkag1* | 0.25 | 0.266 | 0.413 | -0.09 | 0.302 | 0.507 |
| Pathway | *Prkag2* | -0.12 | 0.654 | 0.759 | -0.38 | 0.001 | 0.001 |
| | *Bax* | -0.32 | 0.195 | 0.336 | -0.16 | 0.041 | 0.132 |
| | *Nfkb1* | 0.35 | 0.087 | 0.198 | 0.26 | 0.039 | 0.127 |
| | *Rela* | 0.18 | 0.346 | 0.495 | 0.15 | 0.028 | 0.101 |
| | *Ppargc1a* | 0.74 | 0.030 | 0.110 | -1.68 | 0.003 | 0.018 |
